# Reconstructing the evolutionary history of a functionally diverse gene family reveals complexity at the genetic origins of novelty

**DOI:** 10.1101/583344

**Authors:** Ivan Koludarov, Timothy NW Jackson, Vivek Suranse, Andrea Pozzi, Kartik Sunagar, Alexander S Mikheyev

## Abstract

Gene duplication is associated with the evolution of many novel biological functions at the molecular level. The dominant view, often referred to as “neofunctionalization”, states that duplications precede many novel gene functions by creating functionally redundant copies which are less constrained than singletons. However, numerous alternative models have been formulated, including some in which novel functions emerge prior to duplication. Unfortunately, few studies have reconstructed the evolutionary history of a functionally diverse gene family sufficiently well to differentiate between these models. Here we examined the evolution of the g2 family of phospholipase A2 (EC 3.1.1.4) in the genomes of 93 species from all major lineages of Vertebrata. This family is evolutionarily important and has been co-opted for a diverse range of functions, including innate immunity and venom. The genomic region in which this family is located is remarkably syntenic. This allowed us to reconstruct all duplication events over hundreds of millions of years of evolutionary history using manual annotation of gene clusters, which enabled the discovery of a large number of previously un-annotated genes. Intriguingly, we found that the same ancestral gene in the phospholipase gene cluster independently acquired novel molecular functions in birds, mammals and snake, and all subsequent expansion of the cluster originates from this locus. This suggests that the locus has a deep ancestral propensity for multiplication, likely conferred by a structural arrangement of genomic material (i.e. the “genomic context” of the locus) that dates back at least the amniote MRCA. These results highlight the underlying complexity of gene family evolution, as well as the historical- and context-dependence of gene family evolution.

**Graphical abstract:** 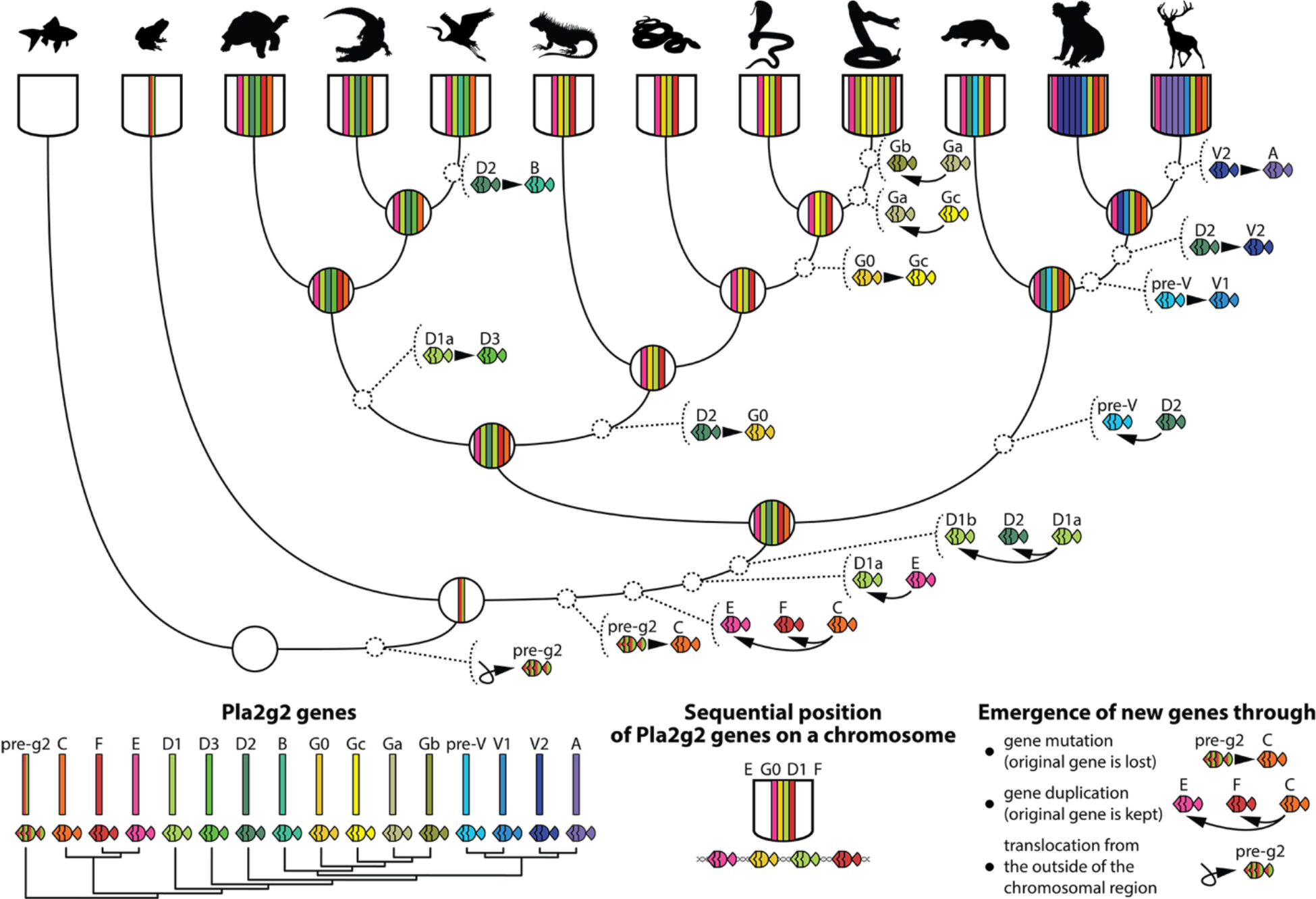

## Introduction

Virtually all studies of gene families have taken a comparative approach focusing on statistical patterns such as copy number variation (CNV) over time. This powerful approach is general and can facilitate the development of evolutionary models to describe the observed patterns, as well as test alternate hypotheses. However, few ancient gene families have been reconstructed in sufficient detail to validate these proposed models. The few exceptions that have been studied, such as the deeply conserved Hox family, are unusual cases, and hard to generalize. As a result, it is not clear whether the models of gene family evolution derived from these studies fit global patterns. Therefore, there is a pressing need to reconstruct gene families in detail, a task that is difficult due to (a) breaks in genomic synteny over large timescales and (b) challenges in assigning orthology within a family after multiple rounds of duplication. Furthermore, understanding the evolution of gene families that have undergone neofunctionalization and positive selection is particularly important as they underlie the origins of phenotypic novelty.

One such family is Phospholipase A2 group 2 (Pla2g2) – a family of enzymes with multiple interaction partners that exhibits CNV in vertebrate genomes. Pla2g2 exhibit structural variation and possess strikingly different functional roles resulting from independent neofunctionalization events in divergent vertebrate lineages – mammalian Pla2g2A genes are important components of innate immunity (Nevalainen 2007; Nevalainen 2008; Birts, Barton, and Wilton 2010), whilst Pla2g2G are a major component of the venom of viperid snakes (Kini 2003). Pla2g2 is also an excellent candidate for comparative genomics research because the cluster is located in a region known to be syntenic across vertebrate genomes – in all these genomes, the region of interest is flanked by OTUD3 and UBXN10 genes (Yamaguchi et al. 2014; Dowell et al. 2016). Although Pla2g2 exists in only 5-6 copies in the genomes of many species, in others the family has undergone considerable expansion associated with the acquisition of novel functions. Notably, the functions associated with gene family expansion are extracellular and “exochemical” – directed towards interaction partners originating outside the body of the producing organism. These functions are likely facilitated (“exapted” – Gould and Vrba 1982) by the ancestral activity of the gene family, hydrolyzing membrane phospholipids, which makes them suitable for deployment as a “weapon” against the cells of other organisms, be they microbes (use in innate immunity) or potential prey/predators (use in venom) (Burke and Dennis 2009).

Comparative genomics is increasingly being heralded as one of the most promising frameworks for understanding the origins of novel functional traits at the molecular level (Drukewitz and von Reumont 2019) as well as organismal evolution and diversification more broadly (Zhang et al. 2014; Dunn and Munro 2016). In the present study, we utilized a comparative approach combining a novel manual genomic annotation technique, phylogenetics, selection rate estimates, and analysis of synteny, to reconstruct the evolutionary history of the Pla2g2 gene family. In the process many new genetic lineages were discovered, necessitating an appraisal and modification of the nomenclature of subclades within this family (see Materials and Methods for the detail). By examining 110 genomic sequences from 93 species across the animal kingdom, we were able to track each duplication event that has occurred since the most recent common ancestor (MRCA) of amniotes, nearly 324 Million Years Ago (Hedges and Kumar 2009). Our analyses identified a number of multiplication events in the gene family’s history, including perhaps the most consequential one, which occurred after the split of Amphibia from Amniota and created the g2 cluster. All extant Pla2g2 genes result from this event. In addition, we demonstrate that a single locus, the same in all lineages, was independently involved in all subsequent acquisitions of novel functionality within the family – birds, snakes and mammals derive proteins with novel functions from the same ancestral gene.

## Results and Discussion

### Structure of Pla2g2 genes and proteins

A typical Pla2g2 gene has 4 exons, the first of which encodes half of the signal peptide, while the three other exons encode what becomes the mature protein. In all figures, we have graphically represented this structure by styling the first exon (signal peptide) as the “tail” and the following three exons (mature protein) as the “body” of each gene (Fig. 1). Our analyses were unable to recover the first exon of some genes and other genes have multiple exons encoding the signal peptide or transmembrane domain (e.g. mammalian g2F – see Fig. 2 for a brief on clades of Pla2g2 and their functions). However, in all the cases the three exons that encode the mature peptide retain the same organization. The number of cysteines (and therefore disulfide bonds) varies from 12 to 15 among the genes, but the most common (and likely plesiotypic) condition is 14 (see SM1 Fig. 13 for consensus primary structures and SM6 for individual sequences).

**Fig. 1.**
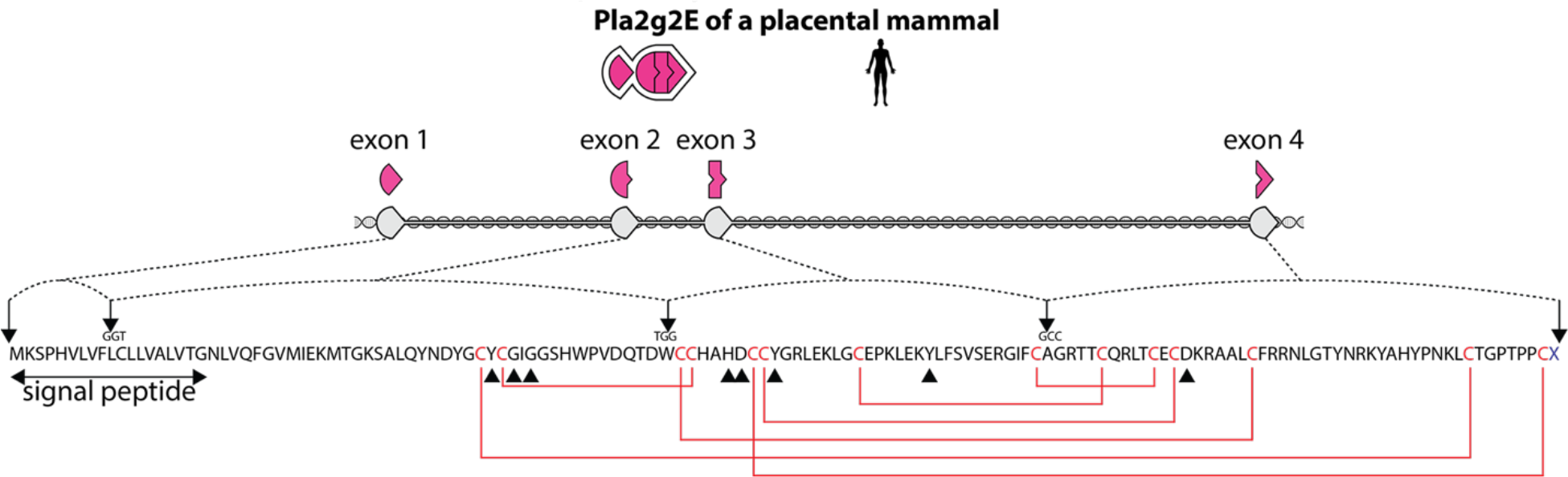
Structure of a typical Pla2g2 gene and the way it corresponds to protein sequence, using *Homo sapiens* g2E as an example (intron lengths are up to scale, cysteines and disulfide bonds are in red, triangles mark catalytic sites, arrows mark the position of the splice sites with respect to codons).

**Fig. 2.**
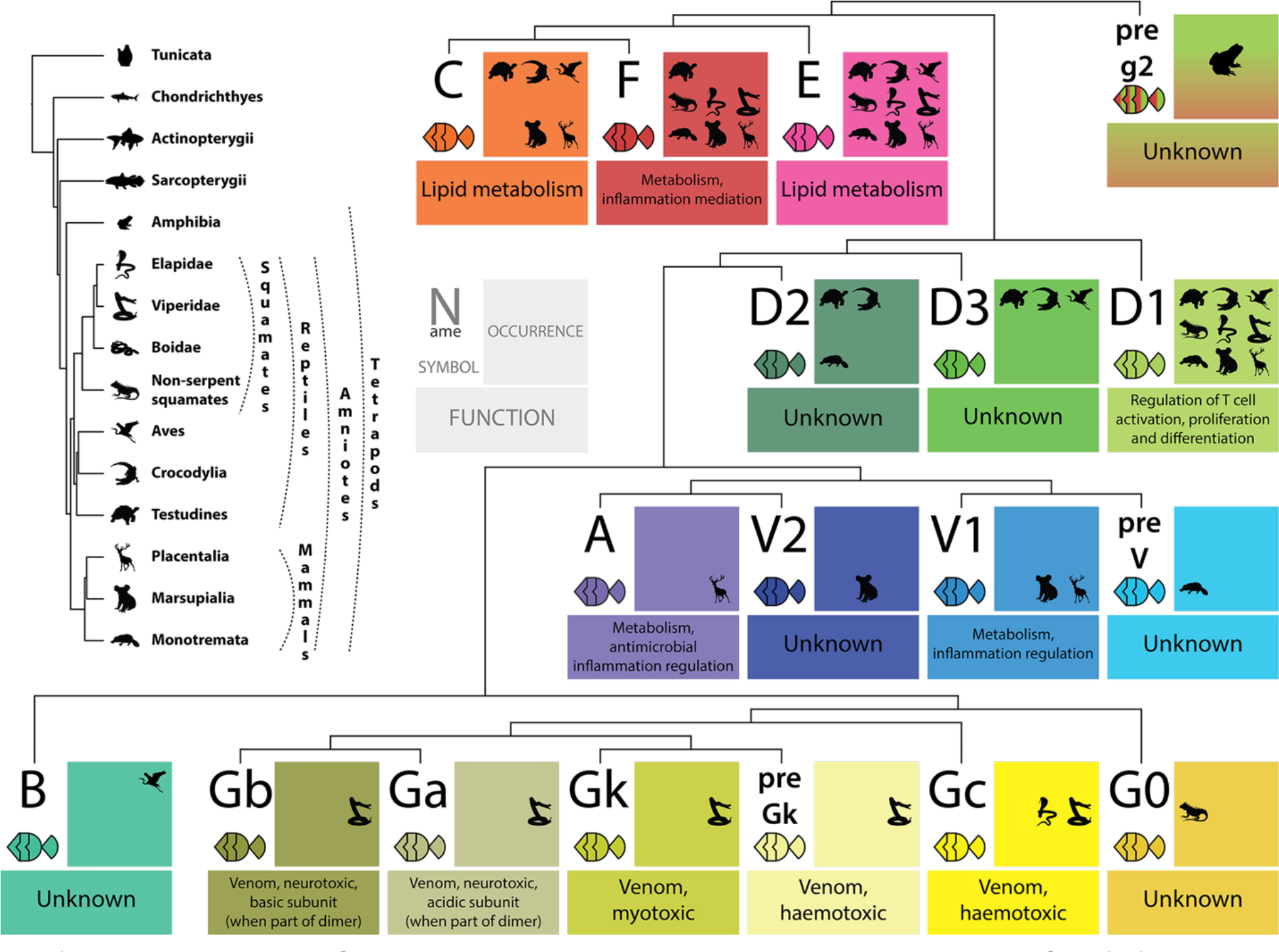
Phylogenetic tree of taxa surveyed in this study and key to color-coding of Pla2g2 clades, along with their phylogenetic relationships, functions and taxonomic presence. Functions as in (Petryszak et al. 2016; Thul et al. 2017; Six and Dennis 2000) and (Fry 2015).

### Pla2g2 gene cluster synteny is conserved in amniotes

We used previously published genomes (see SM2 for the full list) and manual re-annotations to examine the genomic region in which the Pla2g2 gene cluster is located. We discovered remarkable synteny in this region: upstream and downstream regions flanking the Pla2g2 cluster share more than ten genes in almost exactly similar positions across the entire Tetrapoda clade (Fig. 3 and SM1 Fig. 2 and SM1 Fig. 3). This allowed us to reconstruct duplication events spanning 300 million years of the family’s evolutionary history. Interestingly, against this background of conservation, several unrelated species (e.g. *Gecko japonicus, Pelodiscus sinensis*) exhibit substantial rearrangements in this region. In addition to these species-specific rearrangements, the most prominent long-term rearrangement is shared by all squamate reptiles (lizards and snakes), making them more divergent from crocodylians in this region than crocodylians are from humans. Thus, phylogenetic distance is not necessarily an accurate predictor of syntenic conservation in this genomic region.

**Fig. 3.**
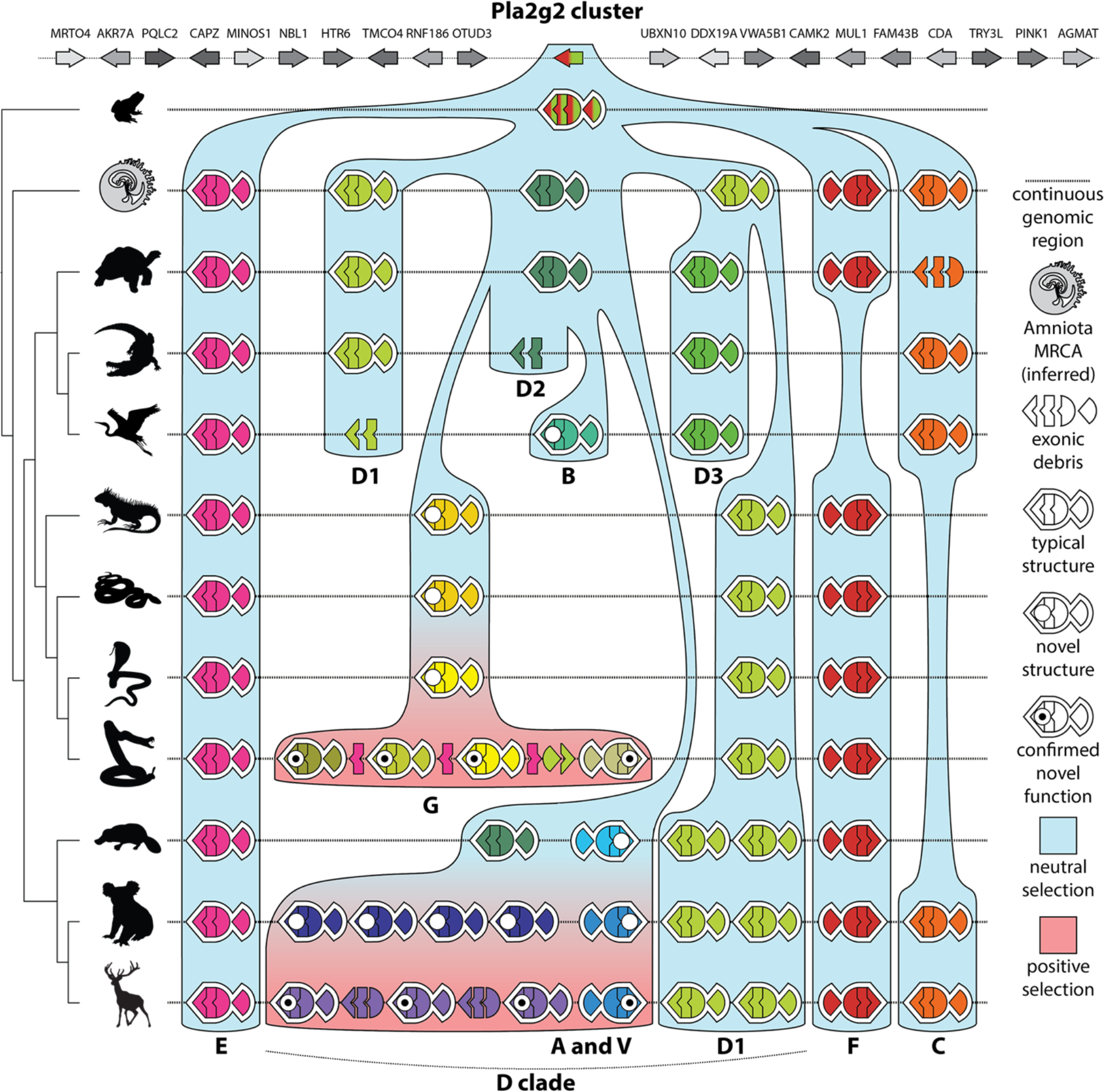
Location of Pla2g2 cluster, representative arrangement of the cluster for each organismal clade; schematic of the phylogenetic relationships among groups; color-coded estimation of selection regimes (blue = negative/neutral; red = positive). Pla2g2 genes are color-coded by group (see Fig. 2 for details).

### Pla2g2 gene clades

All analyzed genes, except those of amphibians, fall into two major clades that diverged from one another some time following the split of Amniota from the amphibians and prior to the evolution of the inferred most recent common ancestor (MRCA) of extant amniotes. These two clades contain Pla2g2s E, F & C; and Pla2g2 D, respectively. Members of the “EFC clade” occupy the flanks of the cluster with g2E genes positioned next to OTUD3 and g2F or g2C positioned next to UBXN10 (Fig. 3). Members of the EFC clade are single-copy genes – E is ubiquitously present, whereas only mammals have functional forms of both F & C (with a few exceptions, e.g. human, in which C is pseudogenized). Birds and crocodylians typically retain C, whereas squamate reptiles retain F. Turtles retain a functional copy of F, but C is typically pseudogenized (see SM1 Fig. 24 for gene maps). The mammalian arrangement, in which both genes are typically functional, likely results from the gene conversion of F from a secretory protein into a transmembrane protein, a novel function unique to this lineage (Thul et al. 2017; Petryszak et al. 2016). Selection analyses indicate that all members of the EFC clade are evolving under the constraint of negative selection (see SM1 Table 1 and SM1 Table 2).

On the other hand, members of the “D clade” occupy the center of the cluster and are involved in all subsequent expansion and neofunctionalization events within the Pla2g2 family (Fig. 3). Both viperid snake g2G toxins and mammalian antimicrobial g2A derive from within this clade.

In our grouping of genes we have tried to preserve previously published Pla2g2 nomenclature created for individual taxa (mostly mammals and vipers, e.g. Six and Dennis 2000), while at the same time reinterpreting this nomenclature in light of the evolutionary history revealed by our analyses (see below for detailed review of Pla2 nomenclature). We have expanded the definition of each Pla2g2 group to make it as concordant with the gene phylogeny as possible and changed some of the names to reflect this phylogeny where it was deemed necessary (Fig. 2).

### Evolution of the Pla2g2 cluster

#### Selection Analyses

To investigate the role of selection in generating sequence diversity within the Pla2g2 family, various Bayesian and maximum-likelihood based site- and branch-site specific models were employed. When grouped into organismal lineages, site-specific analyses indicated that all Pla2g2 (as well as otoconin-22-like genes in all taxa) are evolving under the influence of strong negative selection. The Bayes Empirical Bayes (BEB) analyses failed to identify positively selected sites in all lineages, except one site identified in the avian lineage, while Mixed Effects Model Evolution (MEME) detected 2-7 episodically diversifying sites in all lineages (see SM1 Table 1 and SM1 Table 2). Although the Fast Unconstrained Bayesian AppRoximation (FUBAR) model detected signatures of pervasive positive selection on certain sites in mammals and squamates, it identified a predominant effect of pervasive negative selection on all lineages (SM1 Table 1). These findings may be a consequence of the crucial role played by Pla2g2 in physiological processes, which constrains the accumulation of non-synonymous substitutions that may perturb function. In branch-site specific analyses, we estimated the ω parameter using the two-ratio model test, which indicated the occurrence of positively selected sites in certain clades. Mammalian g2A and g2V exhibited an ω of 2.42, with 1 positively selected site. When all subclasses of squamate reptile g2G were lumped together, the group exhibited an ω of 2.34, with 5 positively selected sites. When the non-toxic form from non-colubroid squamates (g2G0) was removed from this set, analyses returned a higher ω of 2.75 (with the same 5 positively selected sites identified), indicating that the derived toxin forms experience an increased influence of positive selection in comparison to the plesiomorphic gene (SM1 Table 2). The two-ratio model estimated an ω of 2.17 (with 0 positively selected sites) for the incipient toxin g2Gc, however, this analysis failed the likelihood ratio test (LRT). This suggests that this lineage may have experienced positive selection, although the signature was too weak to be confidently identified. Similarly, this test also failed to detect the signatures of positive selection on avian g2B, and the ancient amniote forms g2C, g2E and g2F (SM1 Table 2). Together with the results of the site-selection models and synteny analyses, this highlights the strong evolutionary constraints experienced by the plesiotypic Pla2 gene clusters. Importantly, the combined results highlight the relationship between duplication and positive selection within this gene family, as it is only clades (e.g. g2A and g2G) in which lineage-specific duplications have occurred that exhibit evidence of positive selection (see further discussion below).

#### Reconstructing the ancestral state

Our results allow us to reconstruct the evolutionary history of the Pla2g2 gene family from its origins in an ancient lobe-finned fish (Sarcopterygii) between 400 and 300 million years ago to its diversification in more recent vertebrate lineages. The deeper we go into the evolutionary past, the more we must rely on inference to guide our reconstruction and thus the less credence our conclusions should be given. Nonetheless, the following scenario is suggested by the evidence we have uncovered: after the split of Teleostei and Sarcopterygii, reshuffling introduced a Pla2 gene into the OTUD3-UBXN10 genomic region. This region underwent duplication, either as a large segment or during the whole genome duplications that, according to the 2R hypothesis, occurred early in vertebrate evolution (Van de Peer, Maere, and Meyer 2010). Subsequent to that, a genomic rearrangement resulted in two regions with one Pla2 each – the ancestral g2 gene and Pla2 “otoconin-22-like” gene, the latter being the closest relative to Pla2g2 occurring outside the OTUD3-UBXN10 region. Each of these genes is present in a single copy in *Xenopus* however we were unable to locate any g2 genes *Rana catesbeiana* genome and found only a single g2-like Pla2 gene in *Ambystoma mexicanum*. In contrast to Pla2 “otoconin-22-like” which is always a single-copy gene (if present at all), g2 persisted and presumably gained functional significance in the Amniota clade – by the time of the inferred amniote MRCA it had undergone ancestral expansion to form a cluster of 5-6 genes (Figs 2 & 3).

Based on our analysis, the ancestral g2 gene possessed a structure similar to that of modern g2C. This gene may have triplicated via tandem inversion (TID, SM1 Fig. 29). The evidence for that is the close relationship between the sequences of E, F and C genes that flank the cluster; their direct-reversed-direct position characteristic of the results of TID event; and the presence of a short palindromic sequence upstream of the g2 gene in *Xenopus*, which is necessary to facilitate such an event (Reams and Roth 2015). Alternatively, the gene may have multiplied via two separate tandem duplications.

As mentioned above, while the g2E gene is present in all species studied, there’s a clear taxonomic bias concerning the preservation of the g2F or g2C gene, both of which are present only in the mammals and turtles (with C typically pseudogenized in the latter). In other lineages, sequence similarity between C and F genes may have conferred functional redundancy leading to the elimination of either one or the other.

Early in amniote evolution one of the ancestral EFC genes (possibly the g2E of the Amniota MRCA based on its genomic position) duplicated to create the g2D gene, which in turn spawned at least three additional copies (Fig. 3). This is indicated by the fact that all extant lineages have at least one g2D gene and one or two differentiated D-clade genes. Based on our analysis of synteny, it appears that the amniote MRCA had at least three g2D genes sandwiched between g2E on one side and g2F and g2C on the other. We say “at least”, because there may have been additional copies. The syntenic position and phylogenetic/structural relationships of extant genes indicate that two of the three were members of the plesiotypic g2D1 clade (see Fig. 3).

#### Expansion of D-clade genes is associated with lineage-specific neofunctionalization in snakes and mammals

The major differences in Pla2g2 clusters between different lineages of Amniota concern the evolution of new D-clade genes unique to each taxonomic lineage. All of these genes appear to be descendants of the same ancestral g2D2 gene, which is still present in a plesiomorphic form in crocodiles and turtles (Fig. 3, SM1 Fig.24). The ancestral g2D2 appears to have undergone mutation independently in mammals, birds, and squamate reptiles. These mutated derivations of g2D2 are the ancestors of the g2V (mammalian), g2B (avian), and g2G (squamate reptilian) clades, each of which possesses a unique genomic structure. The avian g2B protein has an N-terminal region unique to this group that sets it apart from almost all other g2. It is always present in only a single copy and its function is unknown. The mammalian and squamate forms are virtually the only ones to have undergone expansion since the ancestral amniote duplication. According to selection analyses, both of these are evolving under the influence of positive selection (SM1 Table 1), however, both of them evolved their unique sequences prior to duplication (Fig. 3).

### Evolution of snake venom Pla2g2 genes

The squamate reptile g2G gene is present in lizards and *Boa constrictor* in a single copy. This gene is the basal member of the clade that contains genes variously labelled as “Gc”, “Gk”, “Ga”, and “Gb” (Dowell et al. 2016). Since it is the plesiotypic (non-toxic) form, we have labelled it g2G0 (Fig. 2). In the ancestor of “advanced snakes” (Colubroidea – i.e. the shared ancestor of cobras and vipers) it underwent structural change whilst remaining a single copy (becoming g2Gc) and this, in addition to the gene family’s ancestral membrane-degrading activity, may have further exapted it for its subsequent functional recruitment into the venom arsenal of vipers. This form, unique to advanced snakes, was recovered from the genomes of the natricid snake *Thermophis baileyi*, the elapid snakes *Notechis scutatus, Ophiophagus hannah*, and *Pseudonaja textilis*, and all viperid species (i.e. all colubroid snakes with genomes of sufficient quality), indicating that it is likely a synapomorphy of colubroid snakes. Readers familiar with snake toxinology should bear in mind that this gene is not the one recruited to the venom arsenal of elapid snakes, which is of a member Pla2 group 1.

Subsequent to the acquisition of a toxic function, a lineage-specific expansion of the gene family occurred within viperid snakes. The first novel isoforms to arise were the g2Ga (acidic) and g2Gb (basic) venom Pla2s (Fig. 4). These speciaised toxin genes are far more abundant in viperid venoms than the plesiotypic g2Gc form from which they are derived (Aird et al. 2017; SM3, SM4). In multiple crotaline (pit viper) lineages, Pla2g2G duplications gave rise to the subunits of heterodimeric neurotoxins (French et al. 2004; Mackessy 2010). The arrangement and orientation of genes and exonic debris indicates that these neurotoxins arose in *Crotalus* and *Sistrurus* independently (Fig. 4; cf. Dowell et al. 2016). This beautiful example of the convergent evolution of a potent toxin is easily explicable as only a single point mutation is required to “unlock cascading exaptations” leading to its formation (Whittington, Mason, and Rokyta 2018). g2Gc (the plesiotypic form) also underwent gene conversion, becoming g2Gck (and intermediate form between g2Gc and g2Gk) in Crotalinae, and subsequently duplicated, giving rise to the non-catalytic myotoxin (g2Gk) (Fig. 4). Whilst the various derived forms mentioned above are abundantly expressed in venoms, expression of the plesiotypical Gc and Gck is apparently suppressed, despite their continued presence in the genomes of viperid snakes surveyed thus far (Aird et al. 2017; Dowell et al. 2016; 2018).

**Fig. 4.**
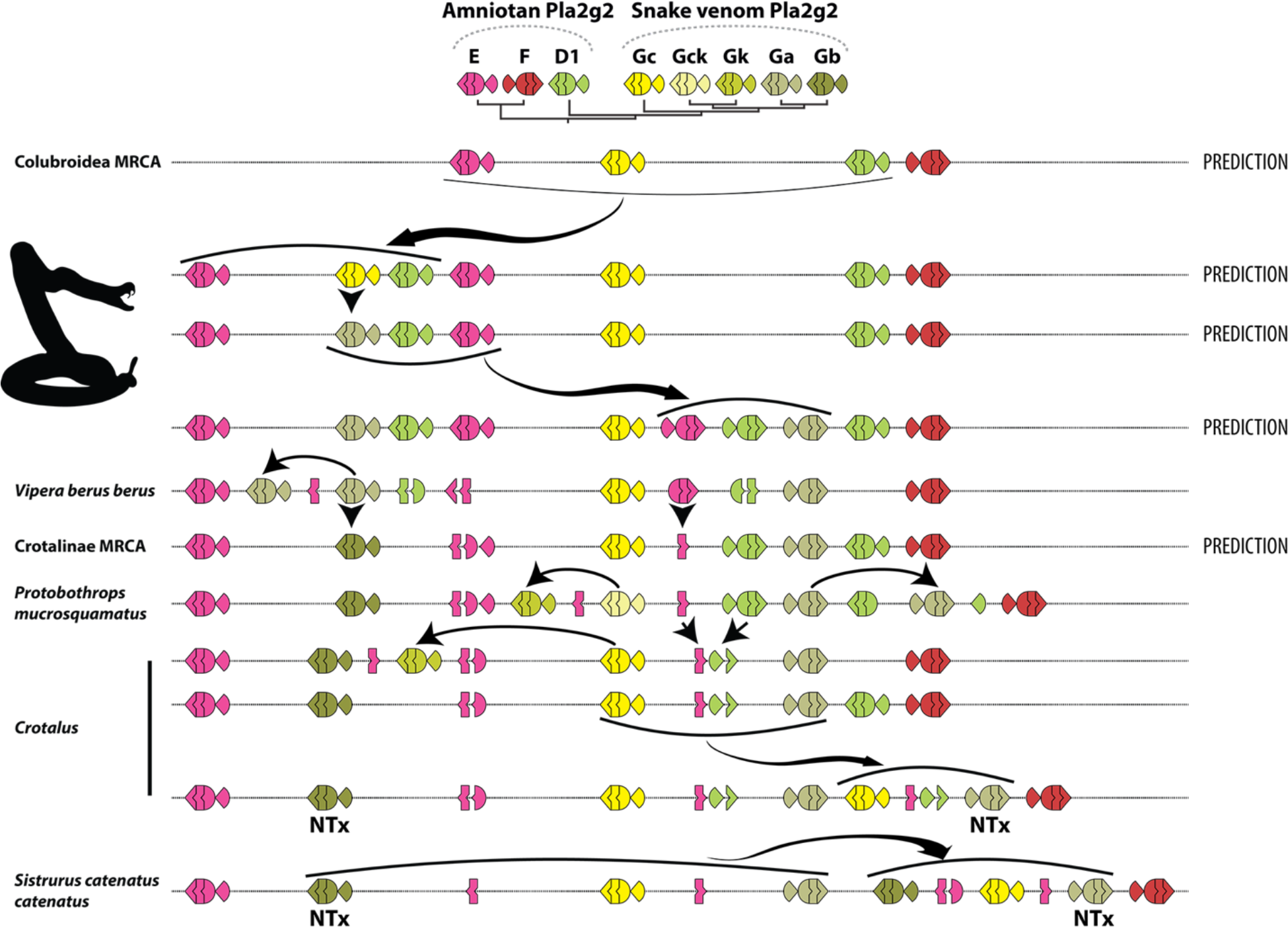
Evolution of Pla2g2 cluster in Viperidae. Note the presence of multiple fragments of “exonic debris” (mainly from g2E and g2D) that make possible the reconstruction of the evolutionary history, including all duplication events, of this genomic region in viperid snakes. Arrows indicate birth and death events. A number of the duplication events (bold arrows) do not involve single genes (unlike g2A of mammals) but rather small groups which are duplicated as units (“cassettes”), typically composed of a g2G gene flanked by parts of g2E and g2D. See SM1 Fig. 24 for precise exonic maps of *Crotalus* and other viperid and colubroid snakes omitted here.

### Mammalian Pla2g2

The g2V clade exhibits a similarly convoluted recent evolutionary history in mammals. An early duplication of g2D produced “pre-g2V” (present in the platypus – Monotremata) and the gene that later became g2V1 (a.k.a. Pla2g5), which is present in both placental (Eutheria) and marsupial (Metatheria) mammals. While g2V1 is always present in a single copy, it likely spawned a form similar to g2V2 of marsupials and then underwent multiple independent expansions (Fig. 3, Fig. 5, Supplementary materials). In marsupials, g2V2-like genes exist in several copies, all of which structurally resemble an intermediate form between g2V1 and g2A of placental mammals, but do not form a monophyletic clade. Group 2A – their counterpart in placental mammals – multiplied independently in different lineages, while being retained as a single copy in some placental mammals (Fig. 5). Most variation (in both structure and number of genes) within placental mammals occurs at the center of the g2A sub-cluster, thus recapitulating the general pattern of Pla2g2 evolution in which expansion takes place at a single central locus (SM1 Fig. 31). It is hard to say how many copies of g2A the eutherian MRCA possessed, given the CNV in both marsupials (Metatheria) and placental mammals (Eutheria).

**Fig. 5.**
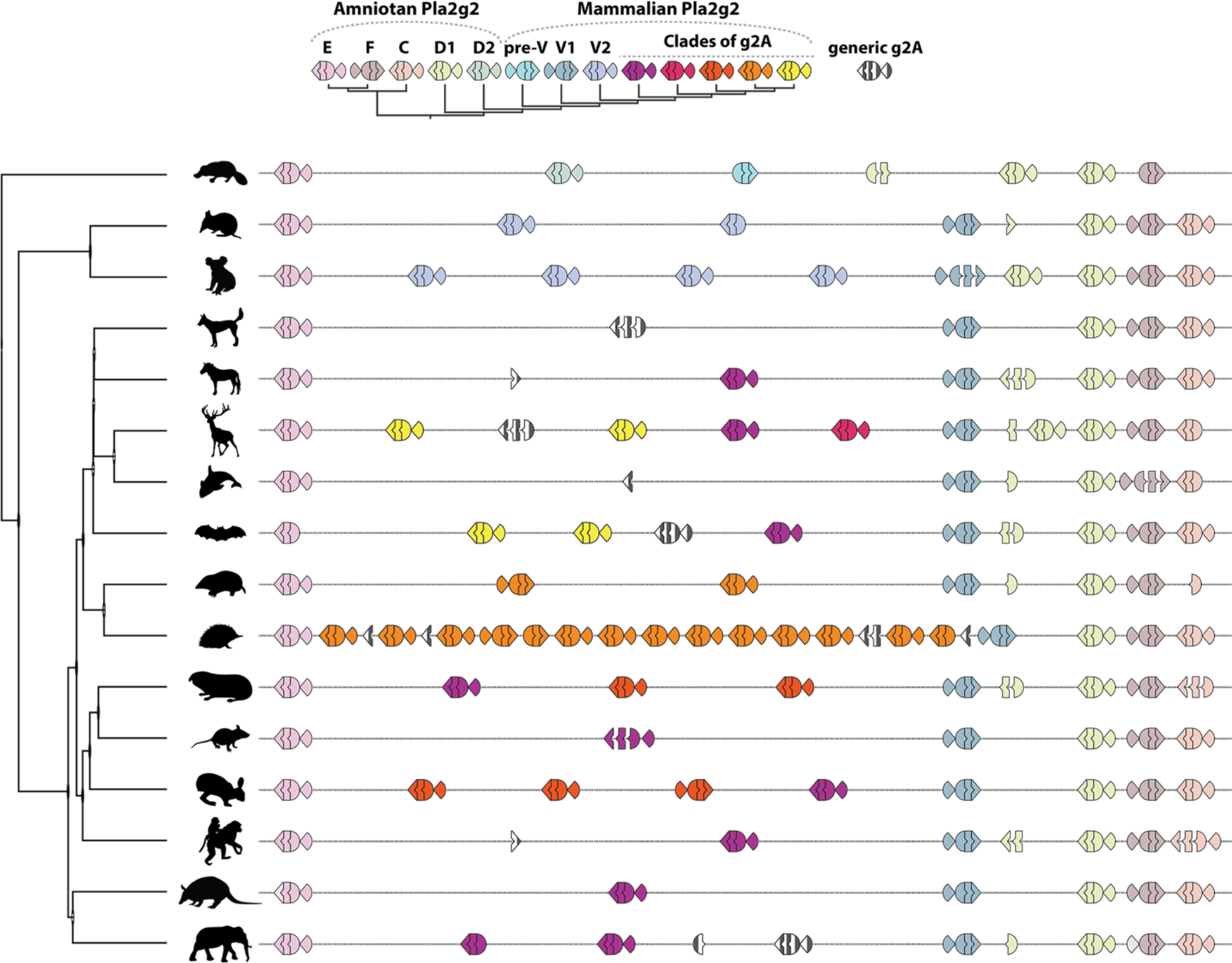
Schematic depiction of expansion and differentiation of mammalian Pla2g2A. Actual complexity of grouping and duplication in placentals is greater, see SM1 Fig. 31 for details. Note that “g2A” forms are confined to placental mammals, although marsupials exhibit distinct patterns of duplication in the same region. Genes and fragments colored green and white belong to the g2A clade, however their relationships within that clade cannot be unambiguously determined. Note that symbols of non-g2A Plag2 forms are partially discolored in comparison to other figures in this study.

Given the aforementioned CNV and the myriad structural forms that have emerged in parallel in distinct taxonomic lineages, group 2A phospholipases appear to be evolving dynamically in placental mammals (Fig. 5). As an extreme example amongst the surveyed taxa, the hedgehog genome contains 14 copies, four of which are successive duplications of a form it shares with moles and shrews. At least one form is shared (deep purple in Fig. 5) amongst (almost) all species that retain a functional g2A gene, suggesting that this gene fulfils an important function. The sole exception to this pattern is the hedgehog, which apparently compensates for the loss of the plesiotypic shared form with its plethora of novel genes. Unfortunately, it is impossible to be precise about the functional roles of products of these genes, given the absence of any functional data or data on tissue-specific expression for any g2A forms from non-model organisms (i.e. >95% of forms). Unlike their single copy g2V1 ancestors, g2A with characterized functions are antimicrobial (Nevalainen, Graham, and Scott 2008). It must be noted that “model” organisms appear anomalous in our data, in that the mouse, human, horse, dog and cat genomes all contain only a single generic form of g2A or lack it entirely. Further research may indicate that distinct forms of g2A should be grouped separately, given that structurally g2A subclades differ as much if not more than venom g2G, and these latter are known to possess strikingly different activities (see discussion above). It is also peculiar that in our dataset only predominantly carnivorous mammals have pseudogenized g2A: the entire Carnivora lineage and *Orcinus orca*. Whether this is functionally significant is an interesting question for future research.

More broadly, based on current data (Fig. 2), it is reasonable to hypothesize that the dynamic evolution of the g2A clade represents innovations built upon a plesiotypic antimicrobial function, which is retained to a varying extent by all members of the group functionally characterized to date. The sister clade of g2A in marsupials – g2V2 – seems to evolve in a similar way, however the paucity of high-quality marsupial genomes and a total lack of activity data for the products of these genes prevents us from reaching any definite conclusion. However, it is clear that the general evolutionary trend of g2V is stable throughout the Mammalia: varying numbers of lineage-specific duplications resulting in novel forms that one can imagine are “fine-tuned” by selection for the particular ecological needs of a particular species. This pattern would be analogous to the evolution of odor receptors and toxin genes. This fascinating evolutionary also story deserves further investigation.

#### A single ancestral gene, g2D2, evolves into a new structural form independently in squamates, birds and mammals

In all three cases (g2B, g2G and g2A/V), these genes evolve new protein structures without prior duplication (Fig. 3). Whilst the avian g2B remains in a single copy, in squamate reptiles and mammals an expansion of the group took place. Interestingly, g2A/V and g2G genes are not just the only Pla2g2 genes to multiply, but also the only ones that show evidence of evolving under the influence of positive selection (SM1 Table 2). This pattern suggests that the acquisition of a novel activity, associated with structural change, perhaps in concert with an appropriate pattern of tissue-specific expression, was the change that facilitated the accumulation of duplicates at this locus. Subsequent to the expansion of these gene networks, the locus became a neofunctionalization hotspot, particularly within viperid snakes and placental mammals.

The observed pattern, in which gene family expansion associated with neofunctionalization occurs at a single locus in distantly related taxa (Fig. 6), suggests that genomic structure inherited from an ancient ancestor common to all these taxa conferred a propensity for mutation and duplication upon this locus. Particular arrangements of genetic material facilitate duplication (Reams and Roth 2015) and thus the propensity for duplication is likely conferred by genomic context. Thus the role of context in the evolution of novel functions is manifold – operating at both the genomic level and at the level of molecular interactions (in which exposure to novel interaction partners provides the opportunity for the emergence of novel functions, cf. Jackson et al., 2019).

**Fig. 6.**
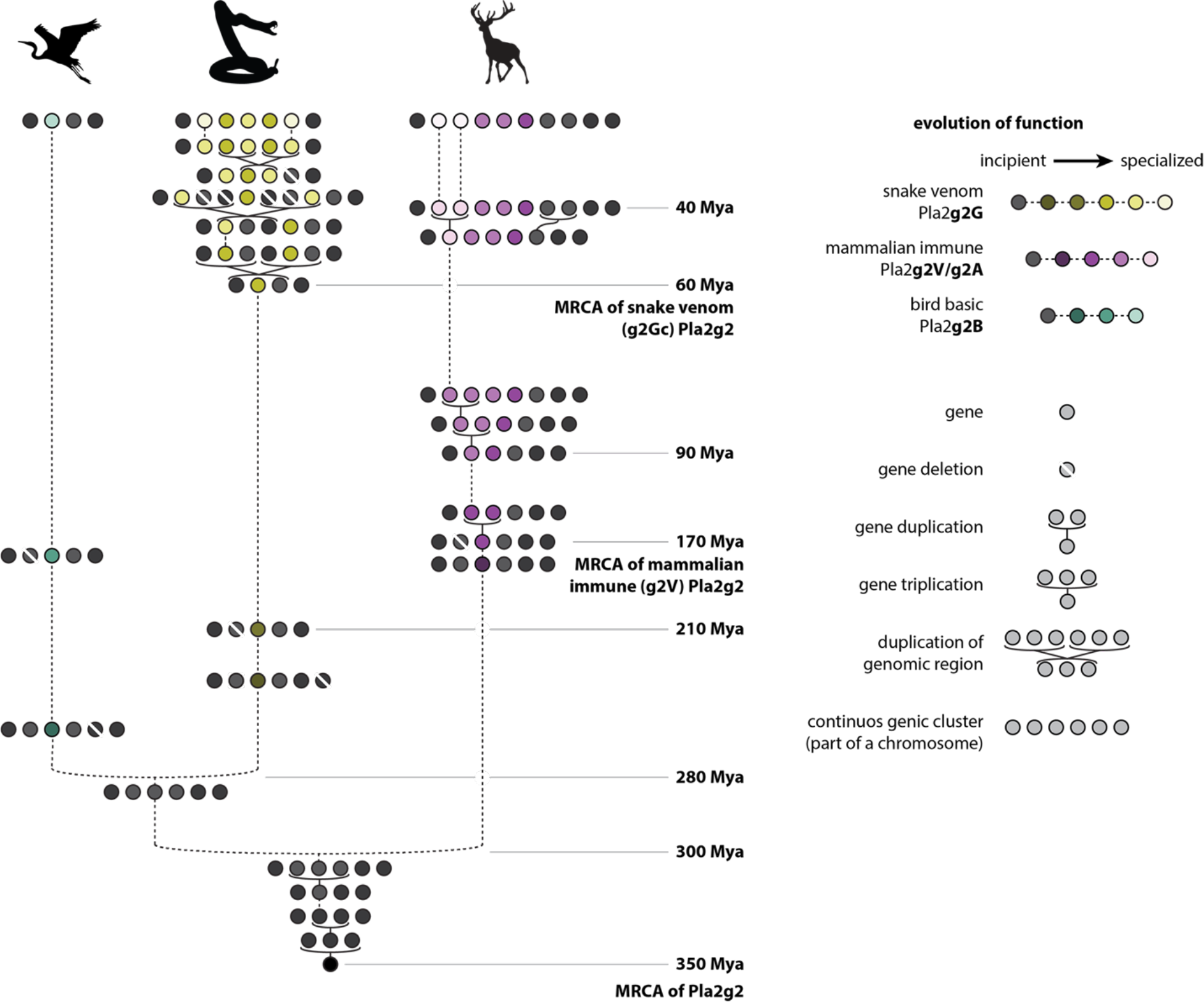
Schematic depiction of the interplay between the emergence of novel functions and duplication in the evolution of Pla2g2. Time estimates are based on data accumulated in TimeTree.org, for details on functions see Fig. 2, for details of synteny see Fig. 3. The figure is based on *Pantholops hodgsonii* for mammals (the pseudogenized g2A is omitted), *Chlamydotis macqueenii* for birds and *Crotalus scutulatus* (form A) for snakes. As discussed in the text, all novel gene forms arise at the center of the cluster, as derivations of g2D2. These novels forms are each given a unique colour. Avian g2B (shades of blue) remains in a single copy in all genomes surveyed. In contrast, in both mammals and squamate reptiles, the gene cluster expands from the central locus, subsequent to the origin of the lineage-specific form g2D2 via mutation. In viperid snakes the evolution is particularly dynamic, with multiplication occurring by both triplication and “duplication” *per se*. The resultant novel forms acquire multiple distinct toxic functions in the venom of viperid and crotaline snakes, an example of classic “neofunctionalization” *sensu* Ohno (see Jackson and Koludarov, *in press*, for further discussion).

## Conclusion

By utilizing an extensive and labour-intensive manual re-annotation method for genomic regions of interest across 110 genomic sequences from 93 species, we have been able to reconstruct the evolutionary history of the Pla2g2 gene family in unprecedented detail. We have thus contributed qualitatively to our knowledge of the evolution of this gene family and developed a method that can be applied to any gene family exhibiting copy number variation and located in a genomic region with a moderate to high level of synteny. We believe that this method, which we have described in detail in the supplementary materials, is complementary with other annotation methods currently being deployed and agree with the assertion made in a recent study of transcriptome assembly pipelines that no single method of assembly and/or annotation is “complete”, or equally applicable to all systems (Venturini et al., 2018). In much the same way as scientific knowledge more generally advances as a consequence of the interaction of a variety of theoretical and experimentally supported arguments and perspectives put forth by a community of enquirers (Popper 1963; Hull 1988; Renn 2020), multiple complementary methods are required to extract the order from complex datasets such as genomes and transcriptomes.

The major theoretical contribution of the paper is the evidence it provides that genomic context is instrumental in conferring a propensity to duplication to loci. The comparative approach furnishes evidence for this hypothesis by demonstrating that all gene family expansion within the Pla2g2 clade takes place at the same locus. Thus, the genomic arrangement (evidenced by the observed synteny) that has shaped the evolution of this gene family is truly ancient, having constrained and facilitated its evolution since the time of the amniote MRCA and likely even earlier.

## Materials and Methods

In recent years the availability of genomic data has increased dramatically. However, our ability to process and interpret this information is yet to catch up with our ability to generate it. Genomic annotation pipelines are still at their best when used to annotate genomes of model organisms or species closely related to them (Wang et al. 2017; Yandell and Ence 2012). Since most organisms don’t fall into this category, much *ab initio* annotation remains challenging.

A potential solution to this problem of *ab initio* annotation is to utilize transcriptomic and proteomic data as a template upon which genomic annotations can be based. A weakness of these homology-based annotation pipelines is that they may attempt to align an entire protein or mRNA sequence with a genomic sequence, a practice that is likely to miss alternative splicing variants or pseudogenes, information concerning which is crucial in evolutionary studies (Danchin et al. 2018; Zhang et al. 2018). In addition, these pipelines have trouble annotating tandem-array duplications, especially when there’s high similarity between the copies (Zallot et al. 2016; Nobre et al. 2016). Assembly of transcriptome libraries is itself challenging and erroneous assemblies may introduce errors that are then perpetuated in genomic annotations based upon them (Venturini et al., 2018).

To address the aforementioned concerns we implemented simple and robust method that aligns genomic sequences with mature exons instead of mRNA, creating an exonic map that then can be further translated into coding sequences (see Materials and Methods and Supplementary Materials for detailed explanation). While this manuscript was in preparation, we used the approach developed for this study to successfully locate and annotate several gene families albeit at a far less ambitious scale: vertebrate NAD glycohydrolases (Koludarov and Aird, 2019), mammalian kallikreins (Casewell et al. 2019) and Indian cobra three finger toxins (Suryamohan et al. 2020). This shows that the approach presented here (for the excruciating detail see Research Notes in SM1) can be useful in studying the evolution of other gene families.

### Locating Pla2g2 cluster

We used published annotations to find genomic sequences that correspond to OTUD3-MUL1 region since those genes were previously shown to be flanking the Pla2g2 cluster in vipers (Chijiwa et al. 2003, Dowell et al. 2016). When no annotations were available, we used the blastn feature of NCBI-BLAST v.2.7.1+ suite to find them (blastn, e-value cutoff of 0.05, default restrictions on word count and gaps), using known sequences as queries. We used the well-annotated and supported by transcriptomic sequencing Protobothrops (Aird at al., 2017) and Crotalus (Dowell et al. 2016; Dowell et al. 2018) genomic scaffolds as starting points and traced single-copy genes contained in that region to their orthologs in non-snake reptiles (lizards, turtles, alligators, birds) as well as mammals. By doing that we established the extreme synteny of the region going as far back as the ancestor of Tetrapoda (see supplementary material SM1 Fig. 1 and SM1 Fig. 2 for genomic maps of the region and comparison between representatives). In all the cases, Pla2g2 cluster was located downstream of OTUD3 gene with MUL1 (squamates) and UBXN10 (all other lineages) genes flanking it on the other side.

### Creating of BLAST database of Pla2g2 exons

All annotated previously annotated exons were extracted from the genomic sequences via “extract” function in Geneious and saved as a fasta file from which initial BLAST database was build from. All genomic sequences in the study were then queried against that database using blastn (blastn, e-value cutoff of 0.05, default restrictions on word count and gaps). BLAST output was formatted into an annotation gff file and imported into Geneious to visualize the results. All exons that had hits against multiple other exons (that is, more than itself and a single closely related species) were exported the same into a new database. All instances when blast revealed previously unannotated exons, they were examined to make sure they complied with the exon structure of Pla2g2 genes that were verified by the transcriptome of a good quality, that is Human, Mouse or snake Pla2g2. Splice cites for each exon were assigned manually based on conserved splice signals for eukaryots. All exons recovered via that method were also put into the new database. Then the entire dataset was queried again using the same dataset tblastx function of NCBI-BLAST suite with e-value cutoff of 0.01. And all the recovered BLAST hits were carefully examined and extracted if they fit the criteria of being a Pla2g2 exon: no frameshift mutations, existing splice sites that do not differ significantly from the known Pla2g2 exons. Then the process was repeated one final time. No new exons were discovered during the last run.

### Creating a dataset for phylogenetic and selection analyses

Since all previously described Pla2g2 genes have 3 exons that encode the mature protein, we considered triplets of those exons (labelled as 2, 3 and 4 respectively) as a separate Pla2g2 gene if they were located in close proximity to each other. The only Pla2g2 group that didn’t conform to the 4 exon rule was mammalian g2F that had 5 exons, with the first two (labelled exon F1 and exon F2) encoding a transmembrane domain, while the other three exons (labelled 2, 3 and 4 for the sake of consistency with other groups) were structurally identical to that of other groups. Coding regions of all genes with intact exons (2,3,4) were extracted and translated for the alignment using Geneious and then aligned with localpair function of MAFFT software v7.305 (Katoh and Standley 2013) with 1000 iterations (--localpair --maxiterate 1000). Alignments were refined by hand using AliView software (Larsson, 2014) to make sure that obviously homologous parts of the molecule (like the cysteine backbone) were aligned properly. The final dataset was trimmed to exclude sequences that might be pseudogenes and included 452 protein sequences

### Phylogenetic analyses of protein sequences

Phylogenetic analysis was performed using ExaBayes v1.5 (Aberer, Kobert, and Stamatakis 2014) software with 10M generations of 4 runs and 4 chains running in parallel (total of 16 chains). The evolutionary model was not specified, which allows software to alternate between different models, until the chains converge on the one that provides the best fit. Final consensus trees were generated with the consense command and the default 25% burn-in. To summarize the run statistics we used postProcParam function and assessed convergence between runs with plots of likelihood and parameter estimates, Effective Sampling Size values higher than 200 and the Potential Scale Reduction Factor lower than 1.1.

FigTree v1.4.3 was used to generate tree figures.

For the construction of species trees we used TimeTree online tool (Kumar et al. 2017), substituting species absent from the database for closely related species of the same genus.

### Analysis of synteny

The overwhelming majority of genes were grouped together on the phylogenetic tree in the way that was totally consistent with their position on the chromosome. The few problematic sequences came from the groups for which only single genome was available – Monotremata and stem anquimorpha that have unique or transitional forms. In those cases we grouped their forms in accordance with the rules we developed for the nomenclature of this gene family (see SM1).

### Selection analyses

The entire dataset that was used for the construction of the phylogenetic trees was used for the analysis of selection. To detect signatures of natural selection and gauge the regime of selection dictating the evolution of various Pla2g2 lineages, site- and branch-site specific maximum likelihood models implemented in CodeML of the PAML (Phylogenetic Analysis by Maximum Likelihood) package were used (Yang 2007), and the omega parameter (ω), or the ratio of non-synonymous to synonymous substitutions, was estimated. To determine the statistical significance of the results obtained from nested models M7 (null model) and M8 (alternate model), the likelihood scores were compared with a likelihood ratio test (LRT). Amino acid sites under the influence of positive diversifying selection were identified using the Bayes Empirical Bayes (BEB) approach in M8 (Yang, Wong, and Nielsen 2005). Data-monkey webserver was used to assess the influence of episodic selection using Mixed Effect Model of Evolution (MEME) and the pervasive effects of diversifying and purifying selection using Fast Unconstrained Bayesian AppRoximation (FUBAR) analysis (Murrell et al. 2012, 2013). For assessing the nature of selection underpinning various Pla2g2 lineages, branch-site specific two ratio model (Yang 1998; Yang and Nielsen 1998) was employed by marking the Pla2g2 lineage suspected to be evolving under positive selection as foreground (ω≥1, alternate model assuming positive selection), and by constraining others as background lineages (ω≤1, null model assuming negative selection or neutral evolution). The likelihood estimates of the null and alternate models were compared with an LRT for determining the significance

## List of supplementary materials

**File SM1.pdf** contains a step by step detailed walkthrough of our methodology and the logic behind it, including (but not by any stretch of imagination limited to):

1. Pla2g2 genomic neighborhood of representative taxonomic clades
2. Convergent evolution of the cysteine arrangement formerly utilized as the defining characteristic of «Group 5» Pla2
3. Results of annotations of *Boa* genome assemblies utilizing different assembly algorithms.
4. Phylogenetic trees of Pla2g2
5. Phylogenetic and syntenic relationships within the placental Pla2g2A clade.
6. Full exonic map for all the species and genomic regions used in this study
7. Alignment of consensus sequences for each of the Pla2g2 group
8. Results of selection analysis

**File SM2.xlsx** contains curated compiled annotation of all genomic sequences used in this study with species names, genomic IDs, references, protein IDs, annotated genes as well as information on their status in previous annotations and key for the naming system used in the datasets.

**File SM3_SM4.xlsx:**

1. SM3: expression data for Pla2s based on previously published studies
2. SM4: expression data for Pm based on Aird 2017

**File SM5_AnnotationsPla2g2.txt** contains concatenated manual annotations made for this study

**File SM6_Pla2g2_aIignment.txt** contains alignment of all the Pla2s used for constructing phylogenetic trees in fasta format

## Acknowledgements

IK would like to thank all the attendees of Venom Evolution, Function and Biomedical Applications Gordon Research Conference 2018 for productive discussions in relation to this study, and Noah L. Dowell from S.B. Carroll’s lab and Michael Broe from H.L. Gibbs’ lab for sharing genomic sequences of *Crotalus* and information on Pla2g2 cluster in *Sistrurus* respectively. IK was partially supported by Japan Society for the Promotion of Science KAKENHI Grant Number 19K16206. KS was supported by funding from the Department for International Development (DFID: grant IAVI/BES/KASU/0002), the Department for Biotechnology, Government of India, through the DBT-IISc Partnership Program, DST-INSPIRE Faculty Award (DST/INSPIRE/04/2017/000071), and DST-FIST (SR/FST/LS-II/2018/233).

## Notes

### Competing Interest Statement

The authors have declared no competing interest.

### Summary of Updates

Edits to the main text.

