## Supplementary Materials 1 for "Reconstructing the evolutionary history of a functionally diverse gene family reveals complexity at the genetic origins of novelty"

### Research notes on the evolution of Phospholipases A2 group 2

**Overview:**

This document is a part of and is intended to be used as a companion piece to an article "Life finds a way: reconstructing the evolutionary history of a functionally diverse gene family reveals complexity at the genetic origins of novelty" by Koludarov et al.

Most of the figures are made as vector graphics and allow for the infitine zoom where needed.

Any suggestions should be forwarded to Koludarov Ivan

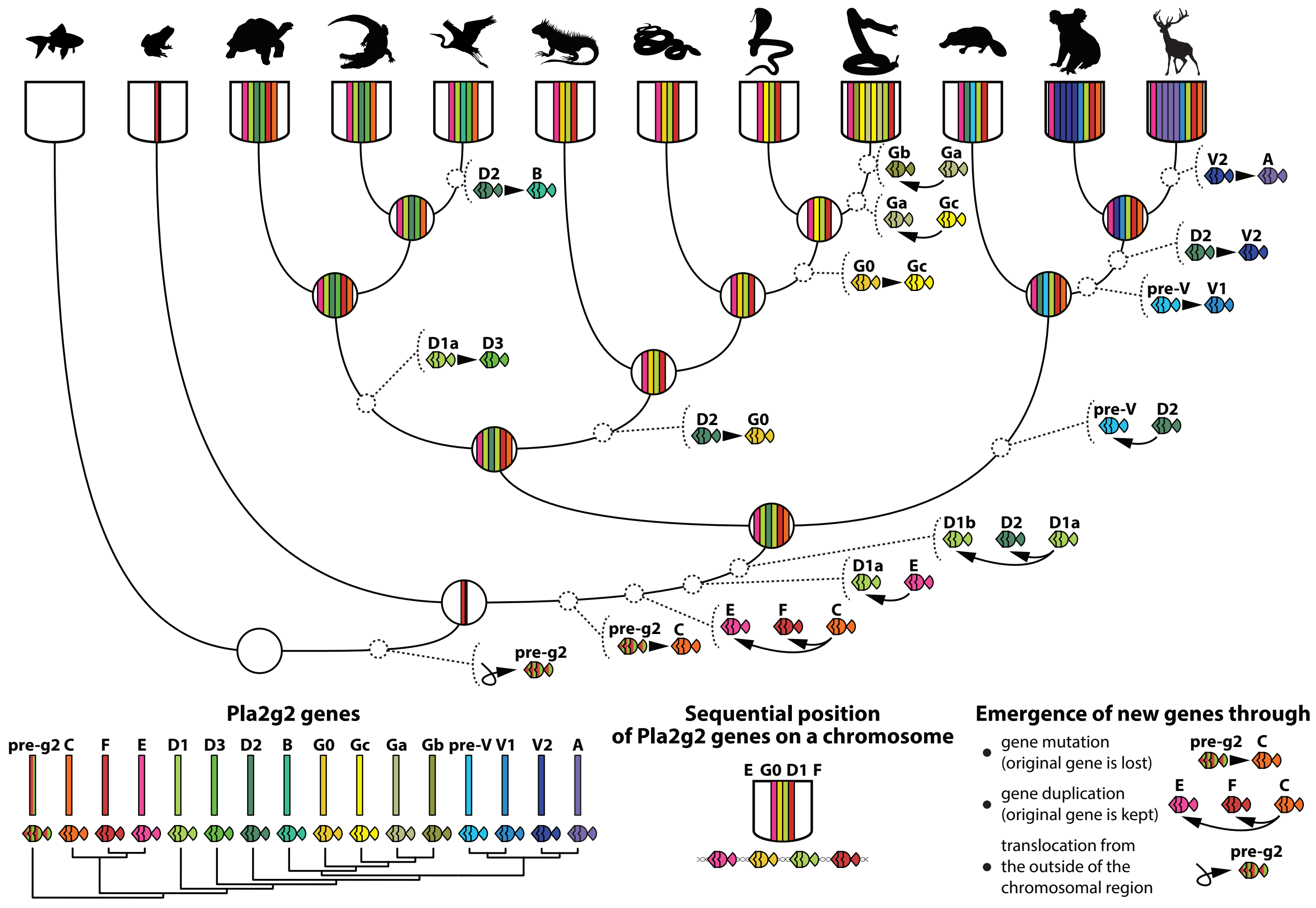

##### *Locating Pla2g2 cluster*

Genomic annotations were downloaded in tabular format for several representatives from each of the major phylogenetic lineages of animals. Taxa were chosen based on the quality of genome assembly, but otherwise randomly. These were transferred into an Excel spreadsheet and the Pla2 cluster was located by searching either for accession numbers from previous papers (Chijiwa et al. 2003, Dowell et al. 2016) or for annotated OTUD3 and MUL1 genes, as these were previously shown to be flanking the Pla2g2 cluster in vipers (Chijiwa et al. 2003, Dowell et al. 2016). After locating genomic scaffolds that contain the Pla2 cluster in representative species, columns that correspond to gene aliases were extracted from the annotation tables of each species and combined to make a bigger table for all representative species (Fig.1). This basic method enabled demonstration of extreme conservation of synteny going as far back as the tetrapod MRCA (Fig. 2 and 3).

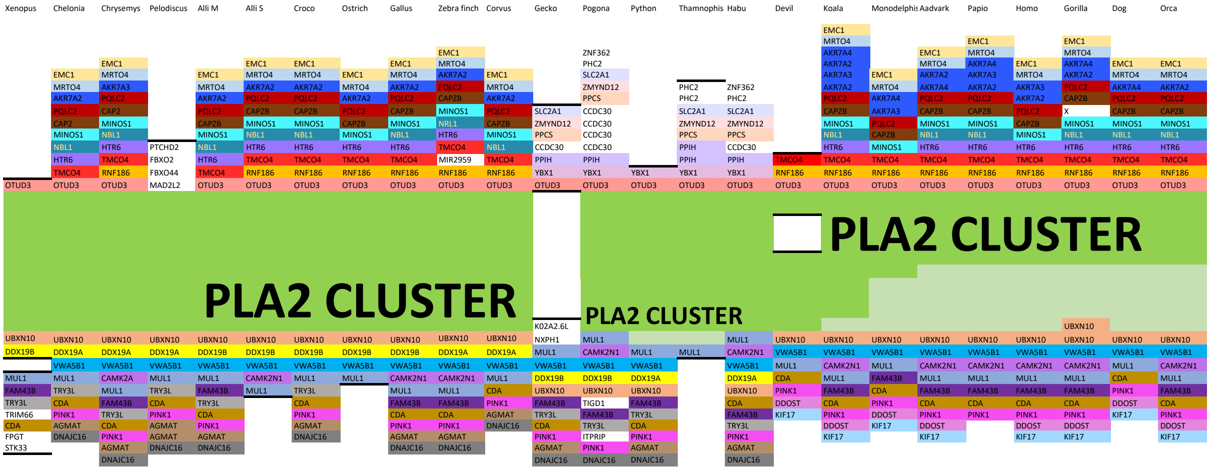

**Fig. 1. Screenshot of an Excel spreadsheet used for initial synteny screening.** Genomic annotations in tab format were loaded into Excel and then columns corresponding to gene aliases for each representative species were combined to form this table. Where no proper gene alias was available, we utilized online nucleotide BLAST to determine the closest annotated homologue and derived gene aliases from there.

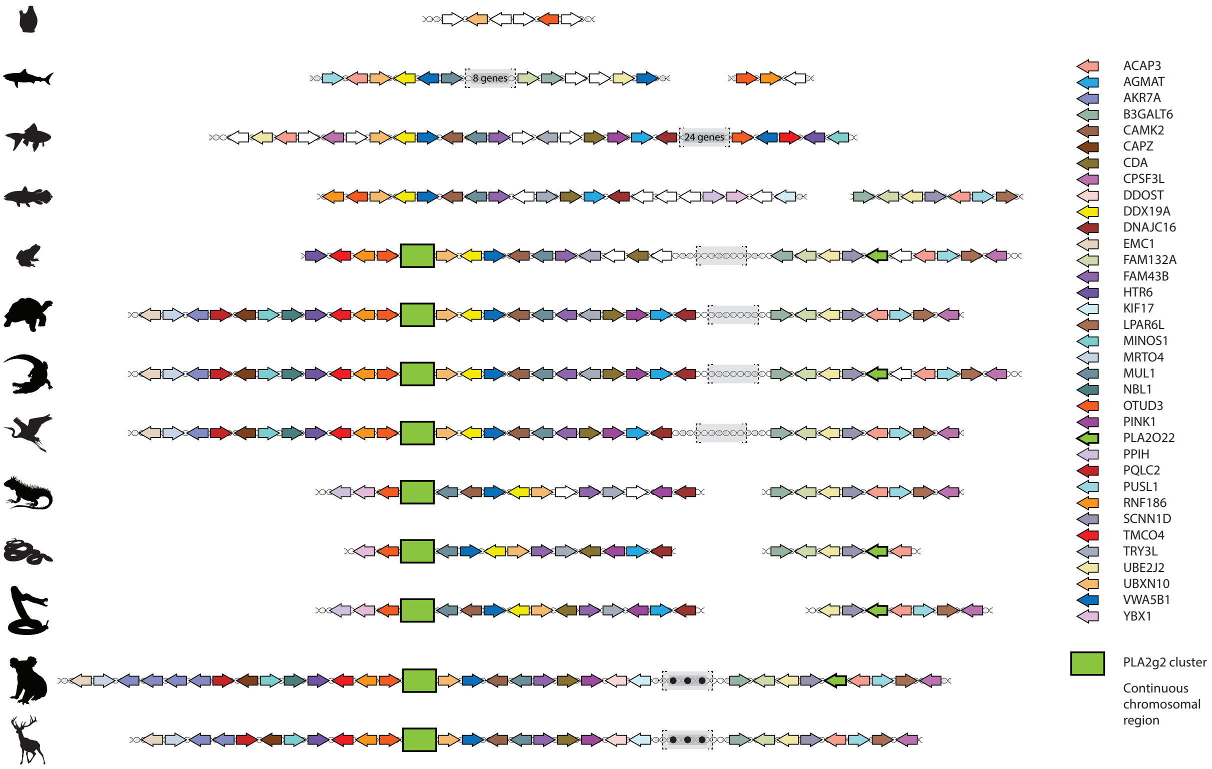

**Fig. 2. Schematic representation of chromosomal synteny surrounding Pla2g2 cluster in vertebrate lineages.** This figure contains the same data as the excel spreadsheet in Figure 1, rendered graphically to aid comprehension.

When no annotations were available, we used the blastn feature of NCBI-BLAST v.2.7.1+ suite to find MUL1; OTUD3; or UBXN10 genes (blastn, e-value cutoff of 0.05, default restrictions on word count and gaps), using all confirmed sequences from representative species to create a search database for each gene. This enabled us to identify the Pla2g2 containing region in previously unannotated genomes. To begin identifying the constituents of the region, we used well-annotated (and supported by transcriptomic sequencing) Protobothrops (Aird at al., 2017) and Crotalus (Dowell et al. 2016; Dowell et al. 2018) genomic scaffolds as starting points and used BLAST (blastn, e-value cutoff of 0.05, default restrictions on word count and gaps) with single-copy genes contained in the region as queries to identify their orthologs in non-snake reptiles (lizards, turtles, alligators, birds) as well as mammals. In all cases, the Pla2g2 cluster was located downstream of the OTUD3 gene with MUL1 (squamates) and UBXN10 (all other lineages) genes flanking it on the other side (Fig. 2 and 3).

This process of reconstructing the Pla2g2 containing region across a wide range of taxa led us to postulate that the ancestral Pla2g2 gene was introduced into the region sometime after the origin of teleost fish and then copied during a major genomic rearrangement that created the OUTD3-UBXN10 region (Fig. 3), within which the Pla2g2 cluster and its closest relative, the otoconin-22-like gene, are located ~40 genes apart from each other (in species where both genes were recovered present).

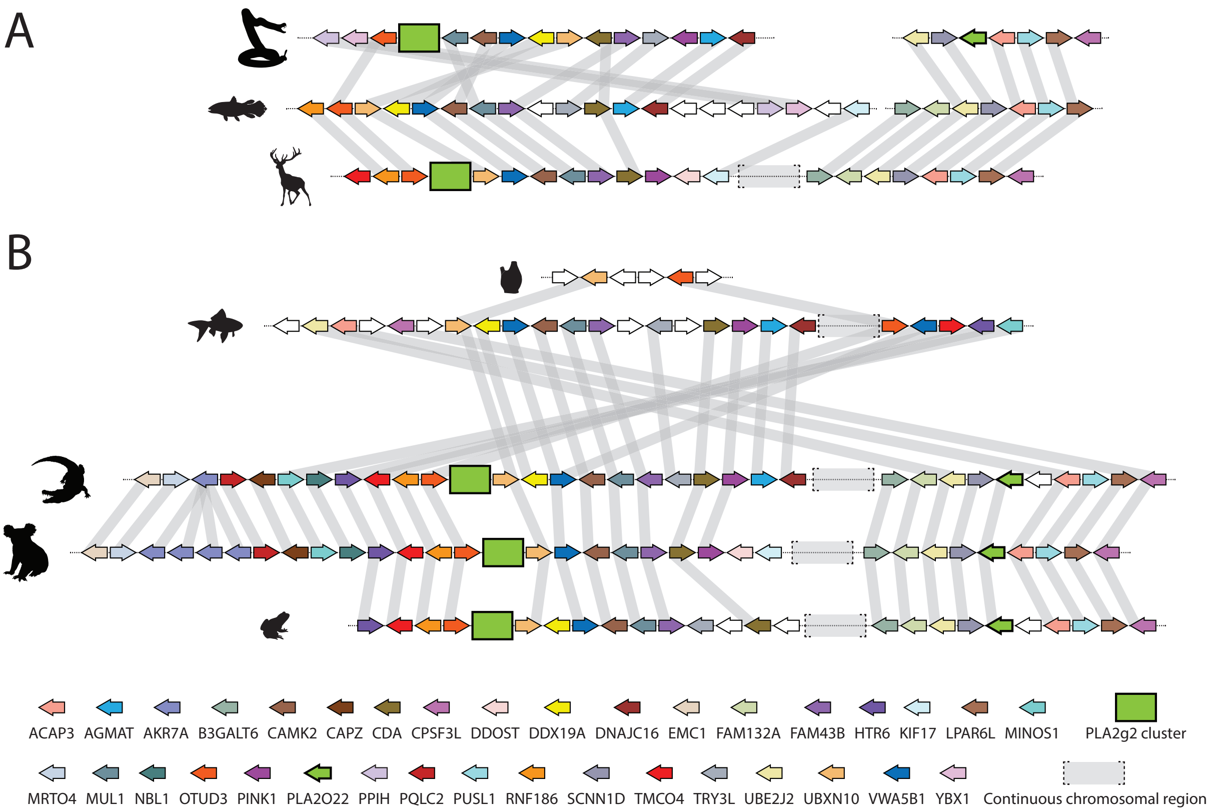

**Fig. 3. Pla2g2 genomic neighborhood and gene orthology as inferred in the present study, grey lines connect orthologous genes.**

#### *Pla2g2 gene structural analysis, gene prediction and annotation*

Initially, all CDS of Pla2g2 genes were extracted according to previously published annotations from representatives of each taxonomic lineage. These annotated genes contained a varying number of exons, making it impossible to align them unambiguously with one another or with encoded proteins, since some of them happen to encode two or more copies of homologous proteins. This results from the fact (as we ascertained during the course of the study) that some of them contained multiple genes (and therefore multiple regions of homology) within a single mis-predicted ORF. We therefore used blastn (blastn, e-value cutoff of 0.05, default restrictions on word count and gaps) to BLAST each individual exon against the database containing all exons. This helped to establish the homology of exons within this sub-set of genes, which revealed that previous annotations of Pla2g2 genes contain differing numbers of homologous exons. During this process, we also uncovered many exons unique to a single species or clade (Fig. 6).

All exons recovered so far were collected into a database for querying (blastn, e-value cutoff of 0.05, default restrictions on word count and gaps) all genomes used in the study to verify the uniqueness

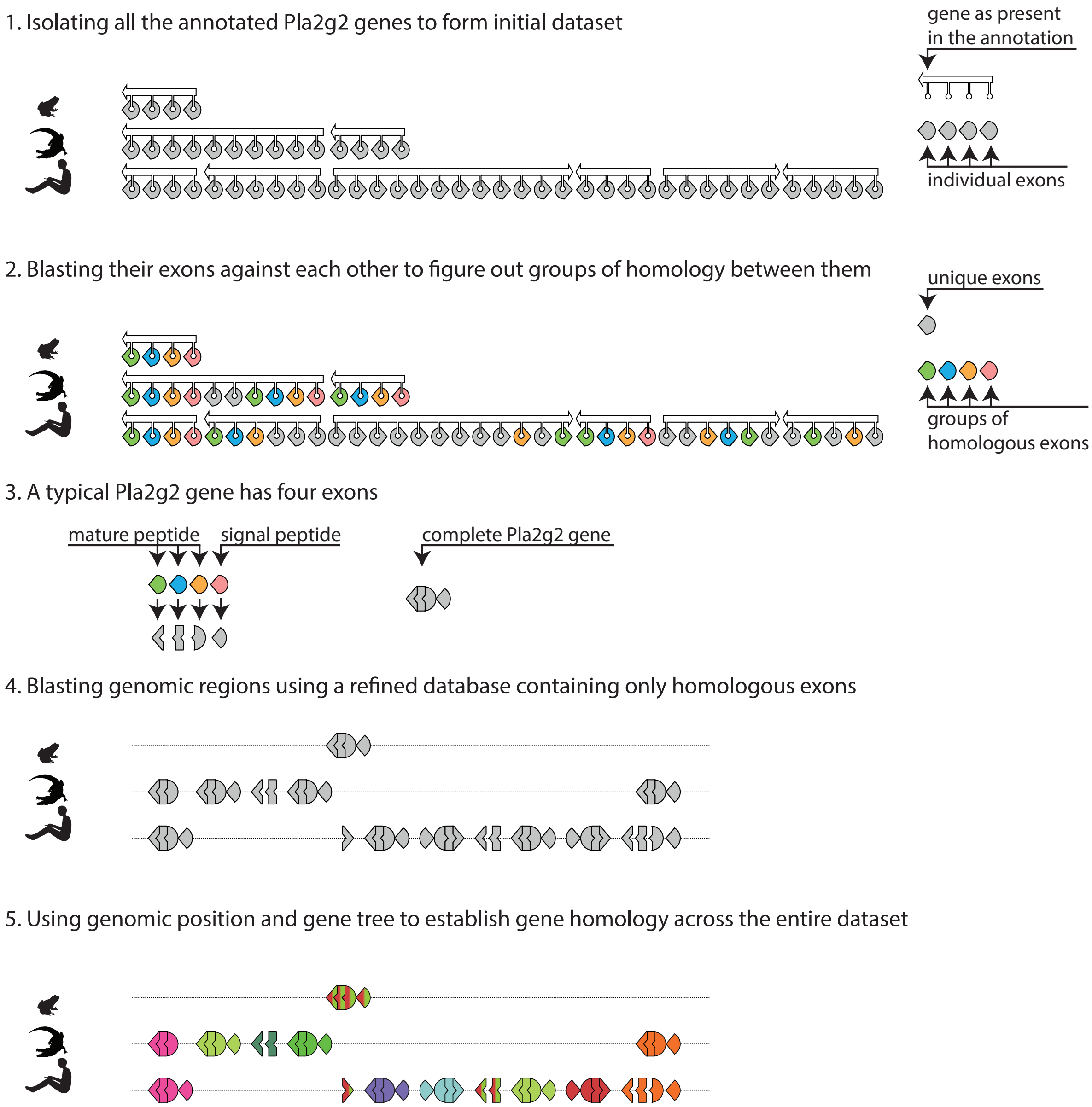

**Fig. 4. Overview of the annotation method depicting three representative species as exemplars.** Previously annotated exons are used to create initial BLAST search database, that is then used to BLAST genomic sequences against. All recovered exons are fed into the database and the process is repeated. All located exons are examined and all possible Pla2g2 genes are annotated.

of singleton exons (i.e. those not belonging to any homology groups).

Cruicially, this process also uncovered exons that were absent from published annotations. We manually annotated these newly discovered exons by adjusting splice cites accordingly to splicing rules and by comparing exon length and splice site position in existing Pla2g2 genes that were confirmed by transcriptomic evidence. All manipulation of exon boundaries and content of annotations, verification of proteins, and translation of CDS into peptides were done within the graphical interface of Geneious v11. Donor-acceptor sites at intron boundaries were assigned manually by locating sequences that conform to the established rules. Each exon's boundaries were checked and confirmed at least three times during the course of the annotation refinement.

The query database was refined by including these newly found sequences and removing singletons and used to search all genomic sequences of interest using the tblastx function of the NCBI-BLAST suite with an e-value cutoff of 0.01.

Boundaries of exons discovered by these searches were established using Geneious v11, relying on previously existing transcriptome-verified exon annotations wherever possible. Variations in exon boundaries existed between the types of exons, groups of Pla2s and between taxonomic lineages and thus these boundaries were scrutinized by eye to ensure that no artefacts were introduced to the data set. We were conservative in our predictions and discarded any annotation that had potential frameshift-inducing mutations or otherwise didn't have the structure of a full Pla2g2 exon. Whether predicted exons are actually transcribed cannot be confirmed without transcriptomic evidence. However, this issue is largely irrelevant for the purposes of our analysis, which is based on sequence homology and order, rather than whether a sequence is transcribed or not. Furthermore, non-transcribed genes can still provide valuable insights into the evolutionary history of the family.

Since all previously described Pla2g2 genes have 3 exons that encode the mature protein, we considered triplets of those exons (labelled as 2, 3 and 4 respectively) as a separate Pla2g2 gene if they were located in close proximity to each other. Exons that encode the N-terminal region (labelled exon 1) of the signal peptide proved much harder to locate and were sometimes present in several copies in a tandem-repeat fashion. This was especially the case in snake genomes with some genes having up to 3 exons of this type, making it impossible to use the full CDS for analysis since it remains impossible to determine which of these exons is present in mRNA without transcript evidence.

The only Pla2g2 group that didn't conform to the 4 exon rule was mammalian g2F, which had 5 exons, with the first two (labelled exon F1 and exon F2) encoding a transmembrane domain, while the other three exons (labelled 2, 3 and 4 for the sake of consistency with other groups) were structurally identical to those of other groups.

This process failed to uncover any Pla2g2 gene with more than 5 exons, indicating that any Pla2g2 gene previously annotated with more than 5 exons is an artefact (with all but mammalian g2F having 4) and no previously annotated Pla2g2 gene with an atypical sequence or set of exons could produce a functional mature Pla2g2. Though it is possible for a Pla2g2 gene (especially a mammalian gene) to have multiple exons preceding the mature sequence, it is highly unlikely for there to exist a Pla2g2 gene with mature peptide coding region containing additional or differently oriented exons than the typical 2, 3 and 4 (as dubbed in this study).

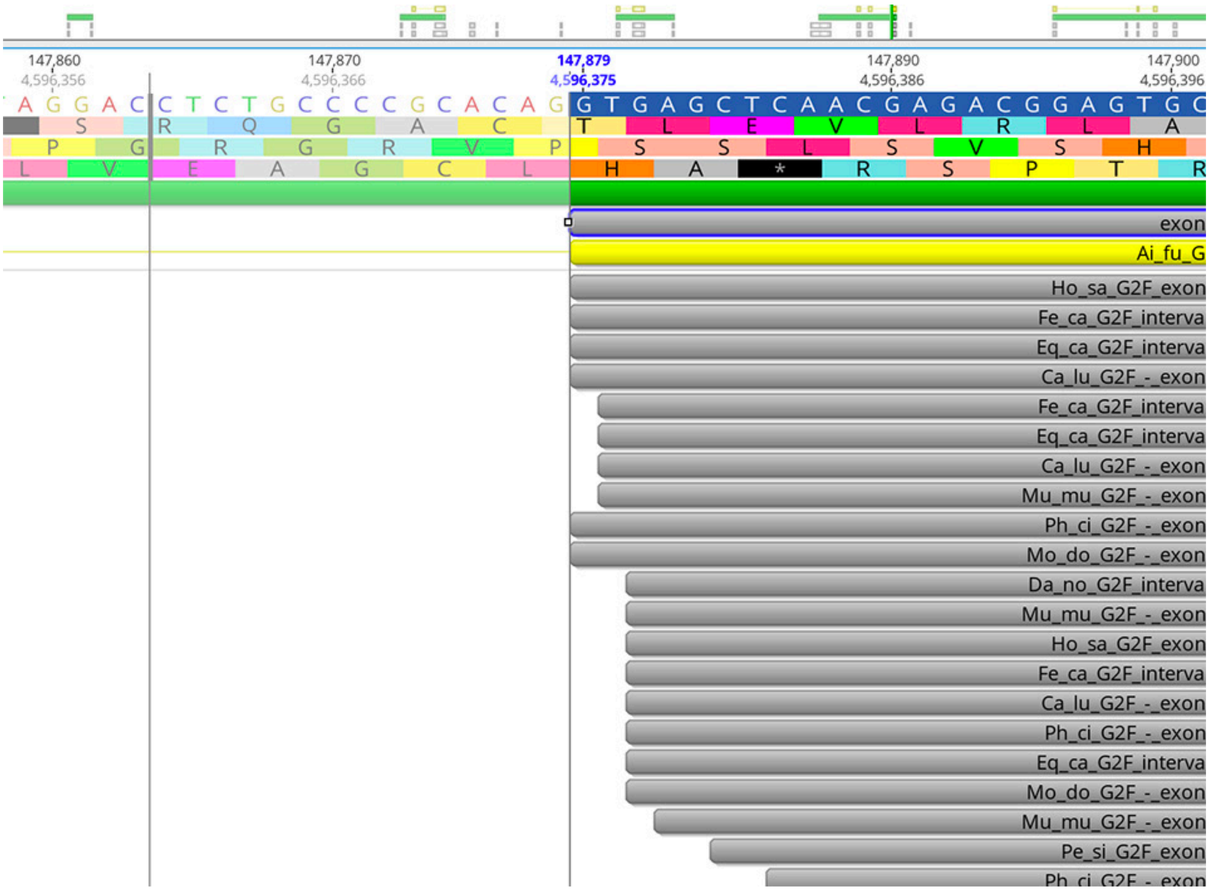

**Fig. 5. Screenshot of Geneious 11 window illustrating the process of adjusting the 5′ splice site of a newly discovered exon and the CDS it is a part of, based on BLAST results and splicing rules.**

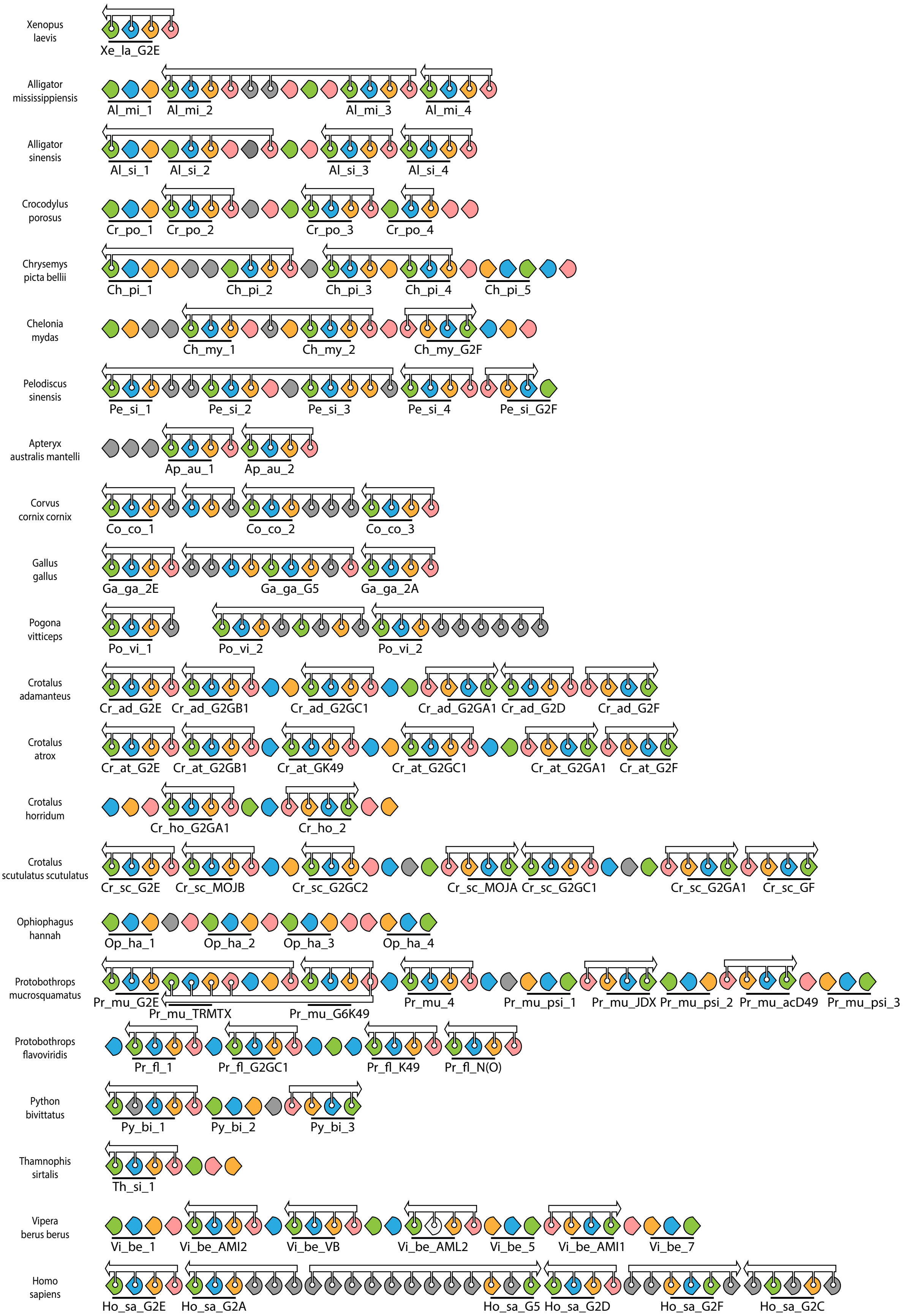

**Fig. 6. Exon homology in representative species.** BLAST results clearly classified exons into five categories: exons homologous to exons 1, 2, 3 and 4 of a typical Pla2g2 gene (color-coded pink, orange, cyan and green respectively) and unique exons with no homology outside of closely related species (grey). White arrows show genes as annotated in public annotations (circa Dec 2018), black lines indicate triplets of exons that form a mature Pla2g2 protein. Gene labels were provisional and used here to indicate the actual genes encoded within this genomic region (vs the annotated genes as shown by white arrows).

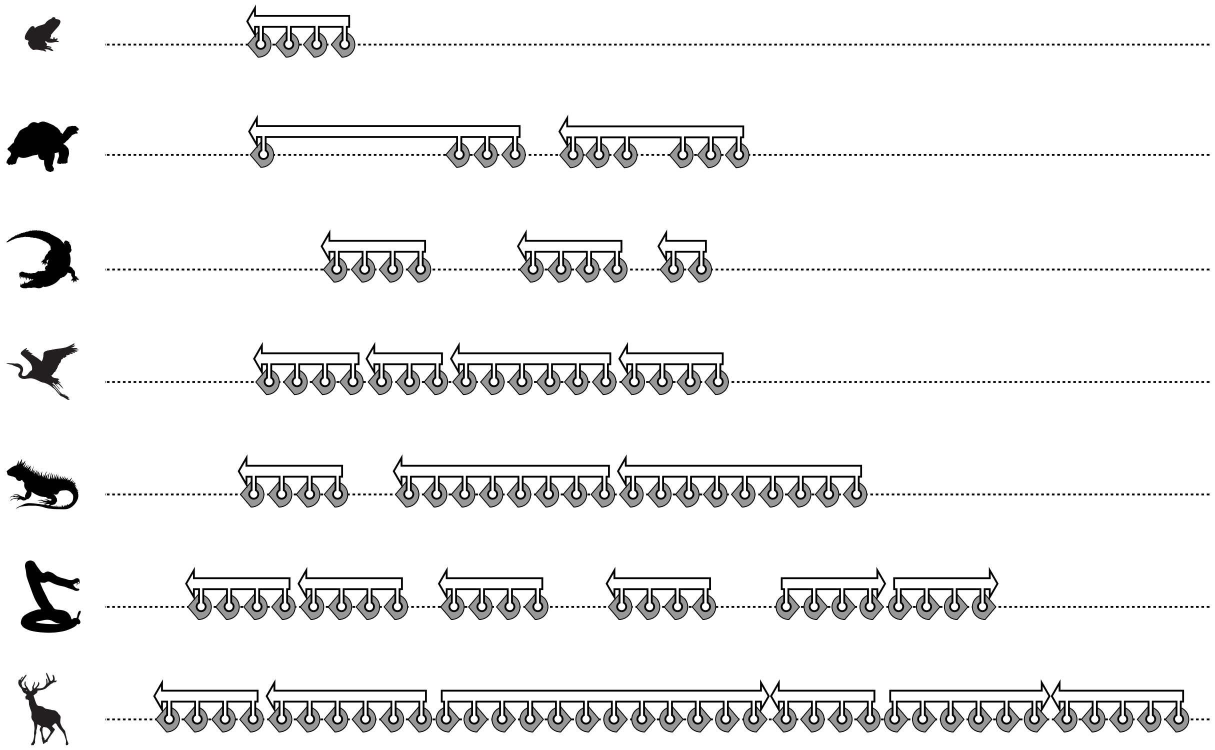

**Fig. 7. Step 1 of establishing homology between Pla2g2 genes: locating Pla2g2 genes as they are annotated in representative species and extracting individual exons belonging to those genes as they are annotated.** The resultant nucleotide database was used as the initial BLAST search database.

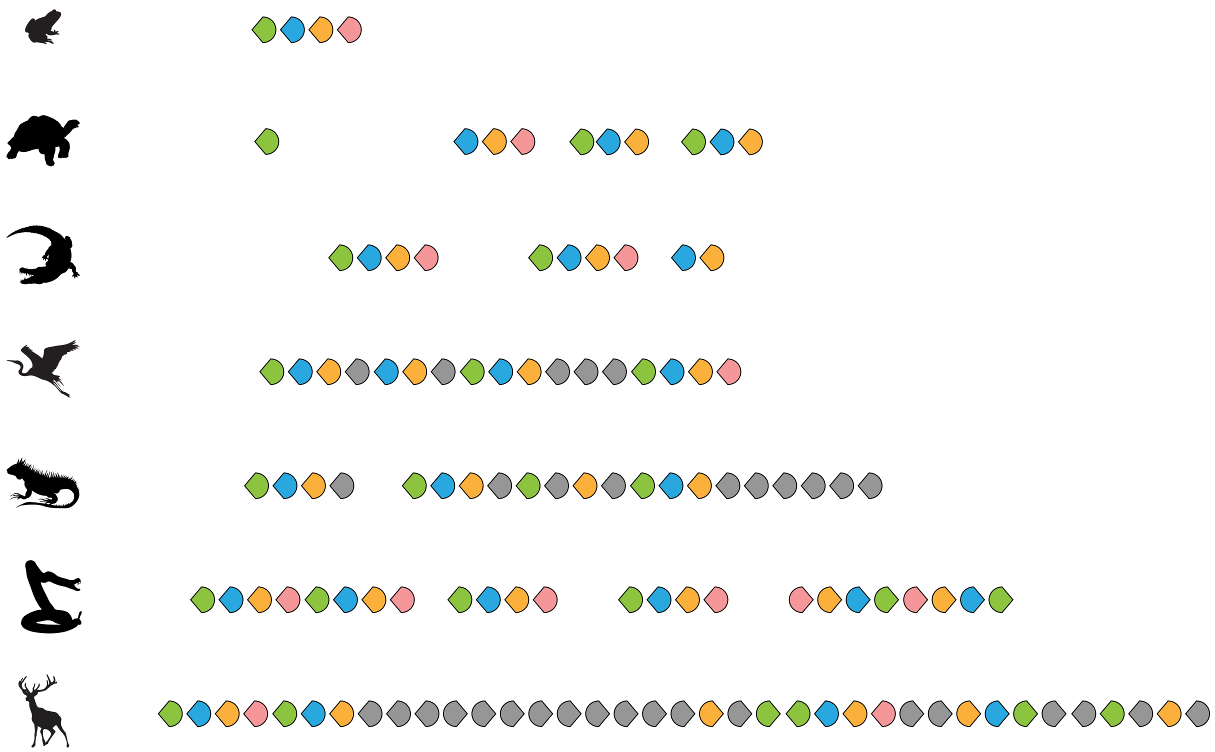

**Fig.8. Step 2 of establishing homology between Pla2g2 genes: establishing groups of homology within the dataset of exons.** This was done by blasting each individual exon against a database containing all exons. Exons homologous to exons 1, 2, 3 and 4 of a typical Pla2g2 gene are color-coded pink, orange, cyan and green respectively while unique exons with no homology outside of closely related species are colored grey.

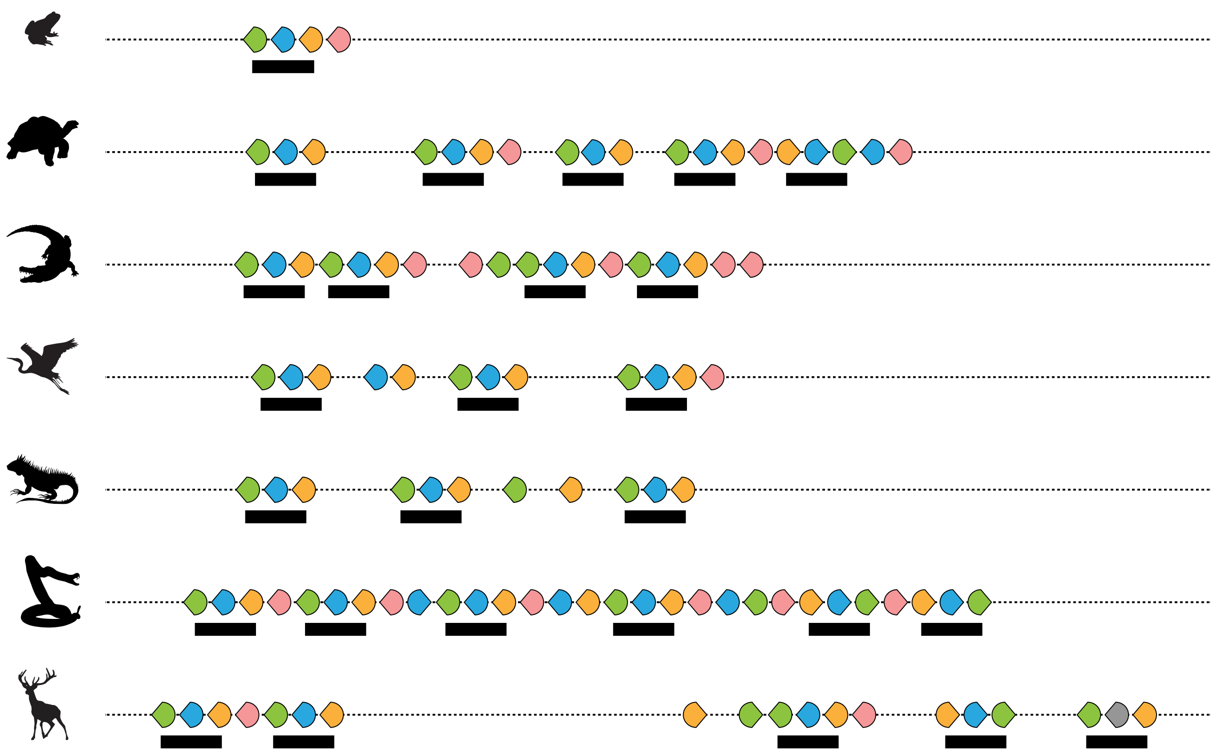

**Fig. 9. Step 3 of establishing homology between Pla2g2 genes: identifying Pla2g2 exons across the entirety of the genomic region.** This was achieved by BLASTing all the genomes used in this study against a database with all extracted exons and grouping them into Pla2g2 genes based on the structure of a typical Pla2g2 gene. Exons homologous to exons 1, 2, 3 and 4 of a typical Pla2g2 gene are color-coded pink, orange, cyan and green respectively, black lines indicate triplets of exons that can produce a mature Pla2g2 protein.

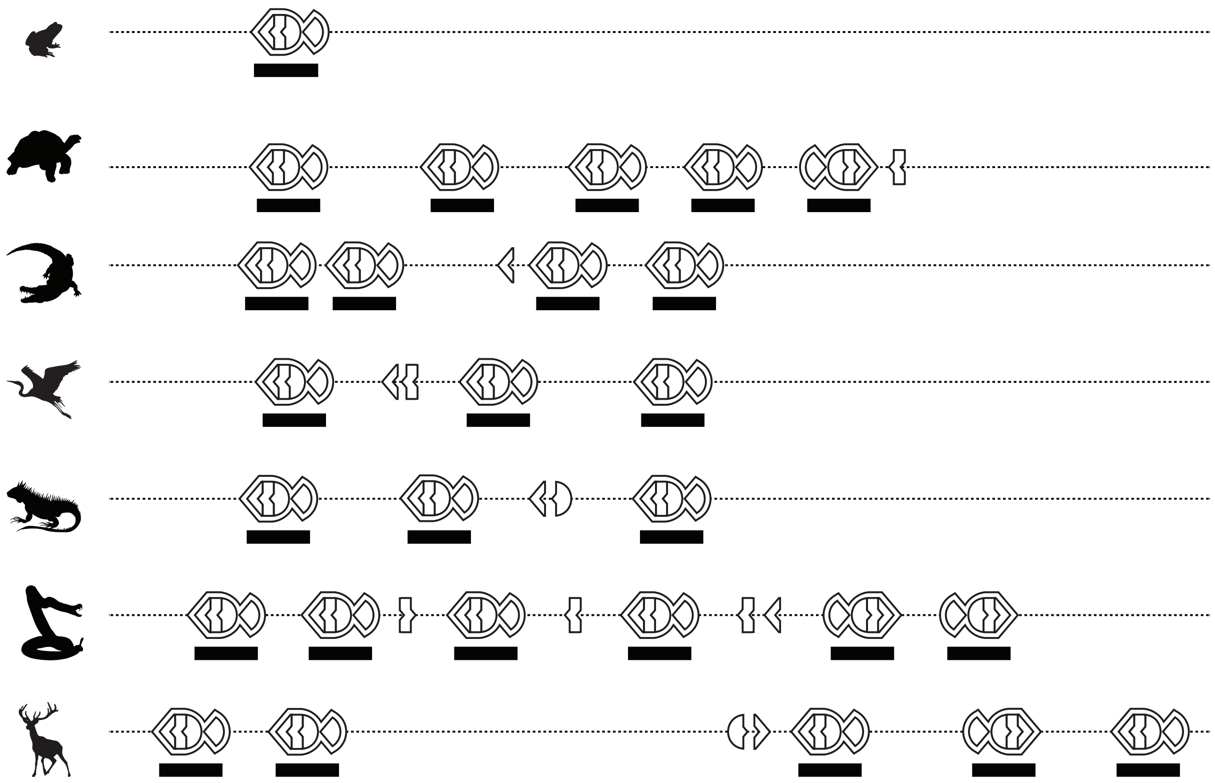

**Fig. 10. Step 4 of establishing homology between Pla2g2 genes: predicting all potential Pla2g2 genes.** Exons that aren't pseudogenized are grouped into Pla2g2 genes if their sequential position in relation to each other allows for a production of a complete Pla2g2 mature protein.

##### *Comparison between manual and automated annotation*

We surveyed 110 individual genomic assemblies in this study, made by different teams using different assembly pipelines (see supplementary material SM2 for full list). Not all of them had gene annotations accompanying them; birds and (especially) mammals typically have much better and more extensive annotations than other vertebrates with exclusion of Xenopus. No non-serpent non-avian reptilian genomic sequence among those surveyed included correctly annotated Pla2g2 genes. This is due to a paucity of transcriptomic data from these taxa and the difficulties associated with using well-studied model organisms such as Mus or Xenopus as templates for the annotation of genomic sequences from highly divergent taxa. All such genomes were either missing genes or had lumped exons of distinct genes together (see Fig. 25 panel A for an example and supplementary material SM2 for the details). Mammalian genomes had much better annotations as a rule, due to several high-quality RNA-seq validated genomic annotations being available, as well the similarities among mammalian genomic arrangements that make them generally comparable to well-studied species such as Homo sapiens. Despite this, most of them were also missing entire genes. It is unknown to what extent this may result from the presence of pseudogenes (which are obviously missed by transcriptome-based annotations), but it showcases that despite advances in annotation technology, an exon-based semi-manual annotation may still be useful to deploy in parallel with currently available bioinformatic pipelines. In future, we intend to explore the possibilities of automating the manual approach we have employed here, though the increasing availability of transcriptomic data may circumvent many of the problems faced by existing automatic homology-based genome annotation. It should be noted, however, that the quality of transcriptome assembly may itself be highly variable, and incorrect assemblies at this level may be perpetuated into genomic annotations based upon them.

Prior to this study, the total number of published annotations available for this genomic region was 71, but this is a misleading number as many of these annotations are incomplete. Some species have only OTUD3 or downstream genes annotated with no Pla2g2 at all, some have only a single correctly annotated Pla2g2E, and others include extensive incorrect annotations. Our study resulted in the net discovery and/or corrected prediction of 135 genes, namely ~30% of all Pla2g2 genes present within the examined genomes.

The average number of Pla2g2 genes differs from clade to clade. Mammals have the highest average (6) along with the best annotations. Though our approach has revealed that many annotations of mammalian genomes are missing entire genes, and most of them are missing pseudogenes, the net gain in that clade was not nearly as significant as in (e.g.) Crocodylia. In the latter clade, of 19 Pla2g2 genes from 4 species only 5 genes had been previously predicted accurately, 5 were entirely absent and 9 were so badly mis-predicted that they might have contaminated the database for that clade resulting in the propagation of the mis-annotation. As a result of such concerns, we resolved to provide

**Table 1. Pla2g2 annotation summary.**

| | | |
|---|---|---|
| Total number of species used in this study | **93** | |
| Total number of genomic sequences used in this study | **110** | |
| Total number of genes annotated in this study | **570** | |
| Total number of genes annotated in previous studies for the same genomic sequences | **395** | |
| Number of previously annotated genes that were verified in this study | **351** | |
| *of those supported with RNAseq data* | 255 | 73 % |
| *of those **not** supported with RNAseq data* | 96 | 27 % |
| Number of previously annotated genes that were found to be erroneous in this study | **44** | |
| *of those supported with RNAseq data* | 30 | 68 % |
| *of those **not** supported with RNAseq data* | 14 | 32 % |

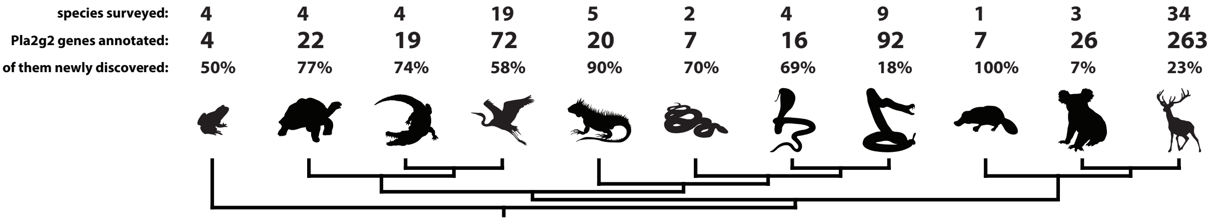

**Fig. 11. Summary of the annotation output performed for this study.** This figure is based on the information provided in SM2 table.

comprehensive information on previous annotations of Pla2g2 in this region and whether or not they were verified by us (see SM2).

In total, we annotated 522 genes and 48 pseudogenes (as well as numerous pieces of "exonic debris") over 110 genomic scaffolds (for annotations see supplementary material SM5, for the summary see SM2). Previously, 395 genes had been annotated on the same scaffolds via different computational pipelines. Of these, 89% (351) were verified by our analyses, while 11% (44) were shown to be erroneous (see supplementary material SM2). Surprisingly, RNA-seq validation of sequences didn't always serve as a good indicator of accurate gene prediction: 73% of correctly predicted genes were verified with RNA-seq coverage VS 68% of incorrectly predicted ones. However, this should not cast doubt yet on the superiority of transcriptome-verified annotations in comparison with ab initio approaches, because of the varying degree of RNA-seq coverage in the genes in question and high similarity between some of the duplicates. Again, the assembly of transcriptomic libraries is itself challenging and as a result these data may be highly variable in accuracy, a variability which may extend to the quality of genomic annotations based upon them (Venturini et al., 2018). These challenges notwithstanding, genes that had close to 100% RNA-seq coverage were generally verified in this study. On the other hand, high similarity between Pla2g2 genes within the same clade (e.g., g2G of snakes or g2A in mammals being the best example) makes unambiguously aligning RNA-seq reads to the genome challenging.

#### *Alternative splicing*

To assess the potential for alternative splicing within this gene family, we surveyed resulting annotations for any set of complete Pla2g2 exons that could facilitate this process. A recent study (Ogawa et al., 2019) failed to find any examples of alternative splicing within toxin-encoding Pla2g2 from viperid snakes and our study corroborates this finding. Exon-skipping, the most common form of alternative splicing in vertebrates (Wang et al., 2015) is unlikely to occur in Pla2g2 as these genes are usually small with short introns, individual genes are located quite far apart from each other, and no additional exons are located nearby that could be incorporated into an alternatively spliced gene product. Similarly, the arrangement of genes makes the use of mutually exclusive exons in alternatively splice gene products unlikely. Other forms of splicing, such as intron-retention, are difficult to rule out, however Pla2g2 are a well-studied gene family at the transcript and protein level, and no exotic forms produced by intron-retention have yet been identified. Indeed, no Pla2g2 gene has been reported to have any kind of alternative splicing, and therefore there is no reason to expect any sophisticated forms of splicing for these genes. We detected only one potential case of alternative splicing within the group (the g2D gene of Bos taurus, see Fig. 24 for the detail), in which functional exons did not conform to the typical gene structure. The resultant isoforms were virtually identical.

Homology of exons that were parts of pseudogenes or weren't part of a complete gene was assigned based on NCBI-BLAST suite best matches. They were taken into account in our overall analysis (Fig. 24) and played an important role in our reconstruction of the evolutionary history of Pla2g2 genes.

#### *Effects of assembly quality*

An Ambystoma mexicanum sequence was excluded from the final analysis, due to our inability to locate the OTUD3 gene and other neighboring genes in the genomic sequence containing it. The lack of these "marker genes" made it impossible to establish the homology of the candidate A. mexicanum Pla2g2 to that of Xenopus. In addition, the gene sequence is divergent enough from other sequences for its placement to be difficult, as our preliminary phylogenetic runs demonstrated. However, it is not different enough for it to be confidently placed in any other group of Pla2s. Therefore it is either a member of Pla2g2 (likely a homolog of Xenopus pre-g2 gene), a member of otoconin-22-like groups of Pla2s (mentioned earlier in the text) or even a member of some distantly related sister-clade of group 2 Pla2s whose history in non-mammalian genomes remains to be uncovered.

It is also worth noting that, as is the case with transcriptome libraries, the quality of the initial genome assembly plays an important role in gene prediction. Many genomes have assembly gaps that may contain exons which may be functional parts of genes. In addition, some methods of assembling a genome are better than others as demonstrated

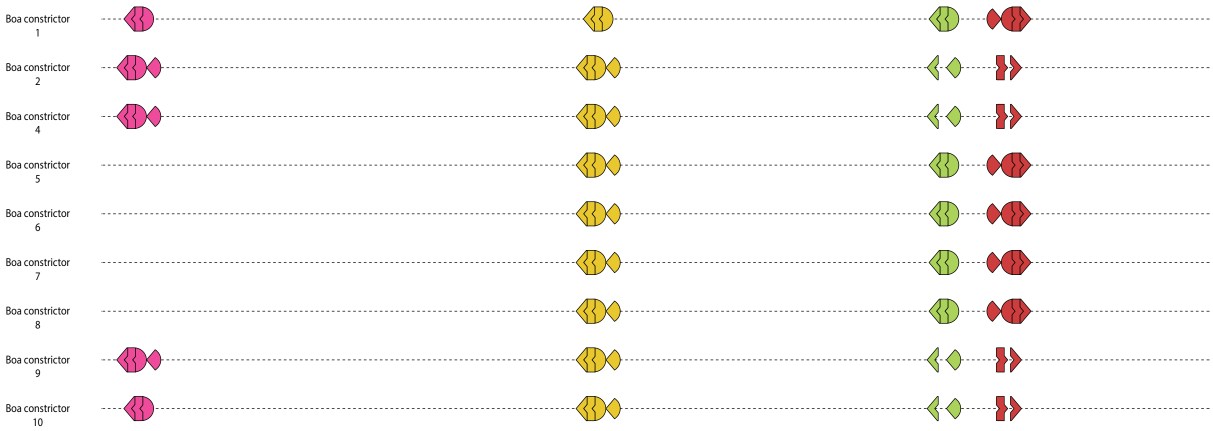

**Fig. 12. Results of *Boa* genome assemblies utilizing different algorithms.** Note the disparities amongst the analyses, including missing exons and entire genes.

for the Boa genome – 10 different assemblies of the same sequencing data resulted in genomes of varying quality [(Bradnam et al. 2013), see Fig. 12 for comparison]. Of the 10, only 2 had complete sequences of all four g2 genes. The absence of a particular gene from a genome in our study may therefore be a consequence of poor assembly.

##### *Alignment*

Individual genes were translated and the mature peptides were aligned using the localpair function of MAFFT software v7.305 (Katoh and Standley 2013) with 1000 iterations (--localpair --maxiterate 1000). Alignments were refined by hand using AliView software (Larsson, 2014) to make sure that obviously homologous parts of the molecule (like the cysteine backbone) were aligned properly. The final dataset was trimmed to exclude sequences that might be pseudogenes and included 452 protein sequences that were clustered into 17 distinct groups based on their sequence similarity, phylogenetic relationship and genomic position with respect to other genes. We created consensus sequences for each group using the online tool WebLogo from UC Berkley.

##### *Phylogenetic analyses of protein sequences*

Phylogenetic analyses were performed using ExaBayes v1.5 (Aberer, Kobert, and Stamatakis 2014) software with 10M generations of 4 runs and 4 chains running in parallel (total of 16 chains). The evolutionary model was not specified, which allows software to alternate between models until the chains converge on the one that provides the best fit. Final consensus trees were generated with the consense command and the default 25% burn-in. To summarize the run statistics we used postProcParam function and assessed convergence between runs with plots of likelihood and parameter estimates, Effective Sampling Size values higher than 200 and the Potential Scale Reduction Factor lower than 1.1.

FigTree v1.4.3 was used to generate tree figures.

Phylogenetic trees were generated for genes grouped by taxonomic clade: Mammals (Platypus, marsupials and placentals), Squamate reptiles (Snakes and Lizards), Aves (birds), Archosauria+turtles (crocodiles, birds and turtles). We didn't include any outgroup sequence in those trees. All of them show branching largely consistent with the tree generated for the entire dataset or representative sequences.

A tree generated utilizing a single representative sequence of each group showed skewed branching, reinforcing the need for good sampling. A tree generated from 5 hand-picked sequences from each Pla2g2 clade (where there were less than five sequences available, all sequences were used instead) resulted in the best node support values of all, and was consistent with the tree generated using the entire dataset.

A graphical overlay of color coding and collapsed branches is presented in Fig. 21. Notice that there are differences between this tree and the final tree of Pla2g2, as 300 million years of gene sequence evolution, including the diversification of highly specialized functional clades (e.g. venom g2G) can distort the results of phylogenetic analyses based on individual gene sequences alone. This results in discrepancies between the results of the Bayesian phylogenetic analyses based upon individual gene sequences and the syntenic analyses based on chromosomal position. When all data, including organismal phylogenetics, is taken into account, all clades have the same internal content, but their relationships to one another are slightly altered.

For the construction of species trees we used in the figures we employed TimeTree online tool (Kumar et al. 2017), substituting species absent from the database for closely related species of the same genus.

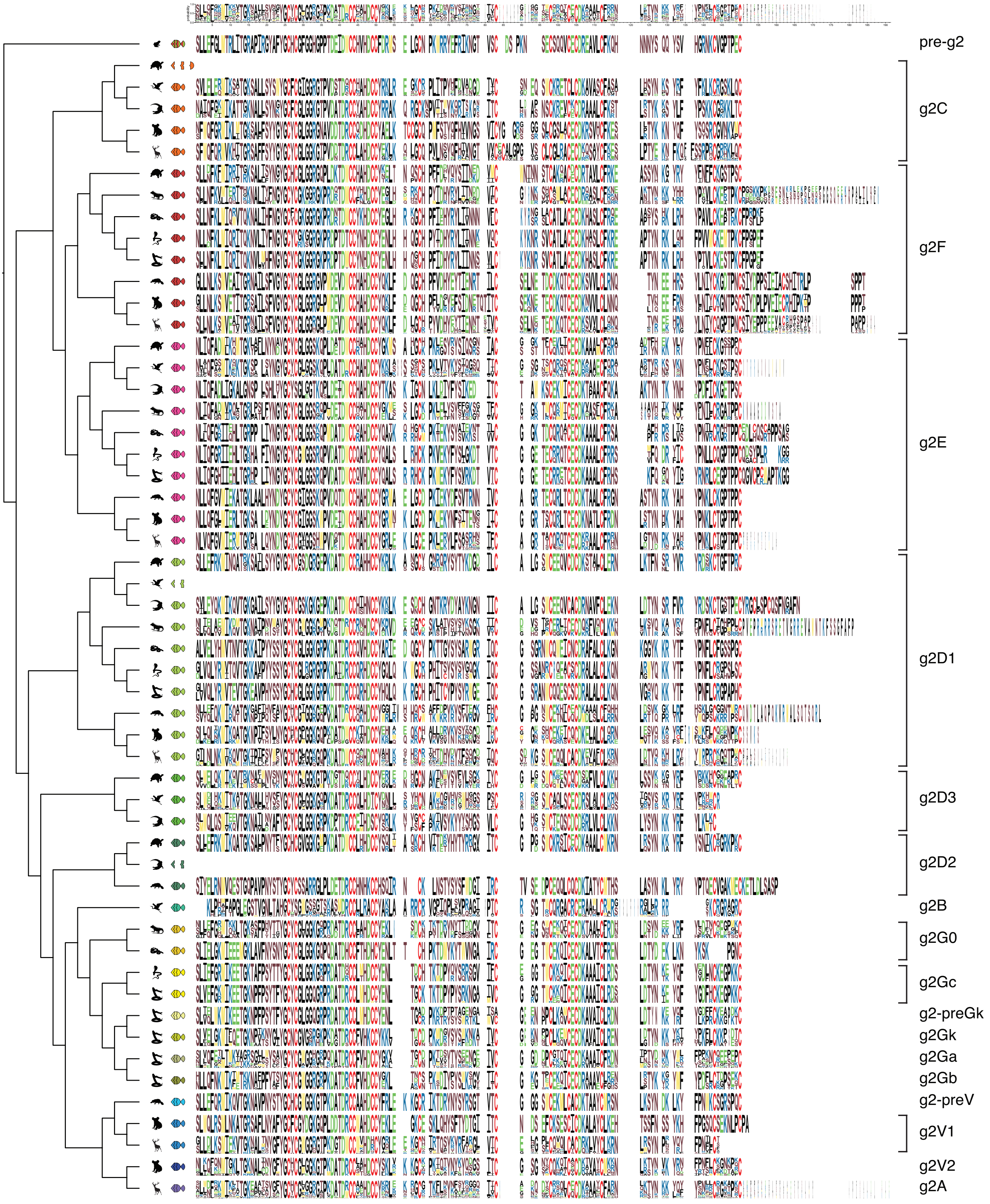

**Fig. 13. Alignment of consensus sequences of Pla2g2 groups.** Consensus sequences were produced in WebLogo online tool.

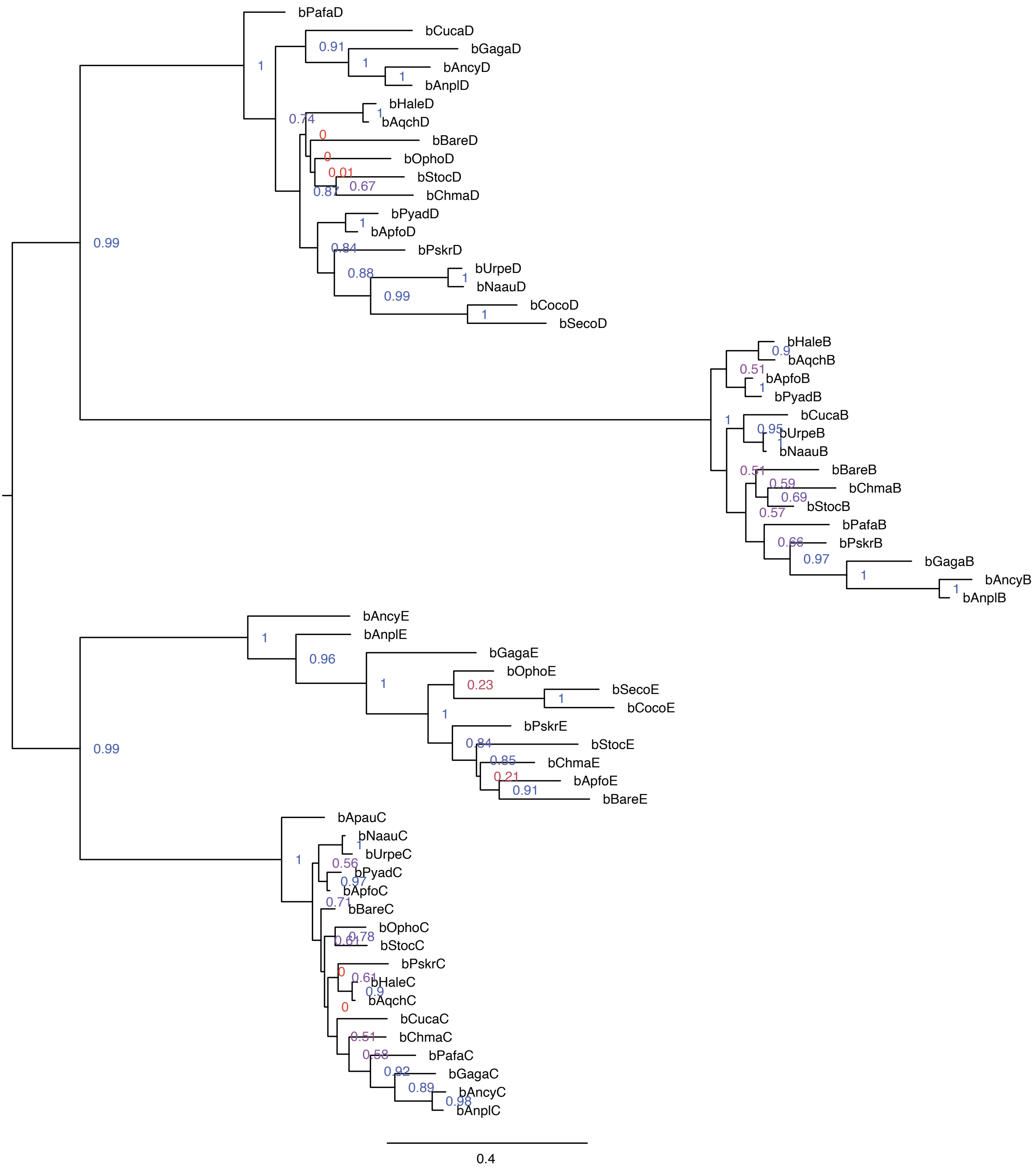

**Fig. 14. Phylogenetic tree of Pla2g2 genes of Aves.** The tree is rerooted on the split between D-clade and EFC-clade.

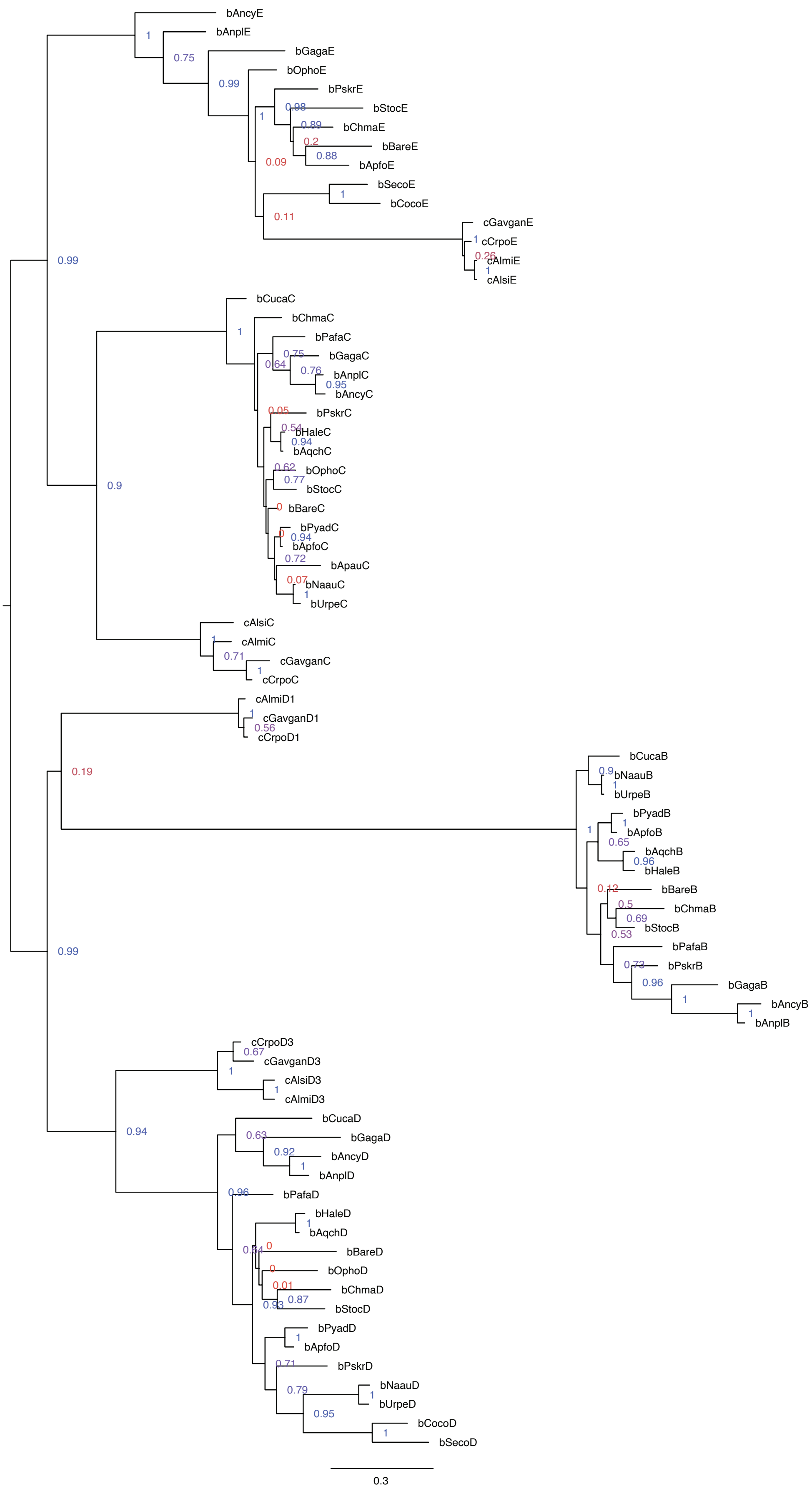

**Fig. 15. Phylogenetic tree of Pla2g2 genes of Archosauria.** The tree is rerooted on the split between D-clade and EFC-clade.

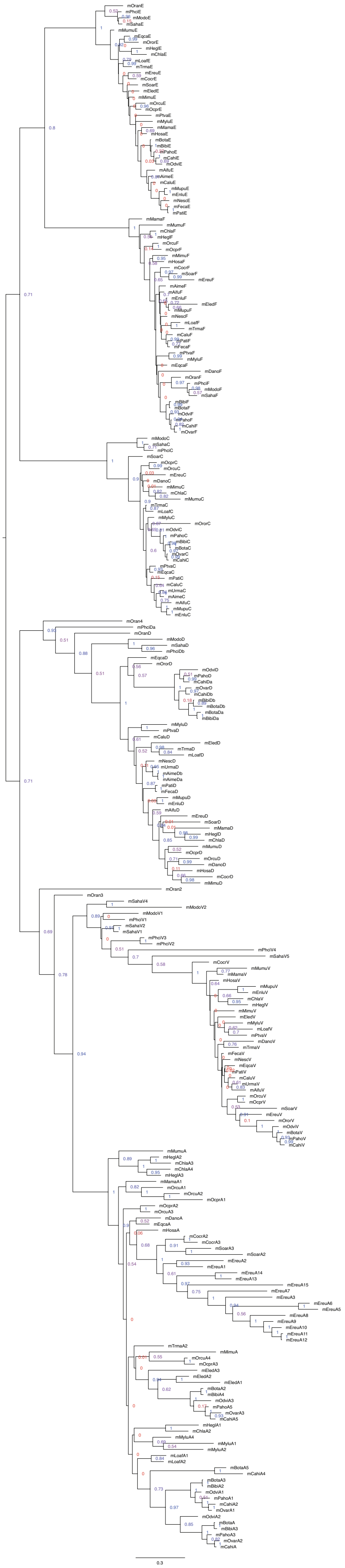

**Fig. 16. Phylogenetic tree of Pla2g2 genes of Mammalia.** The tree is rerooted on the split between D-clade and EFC-clade.

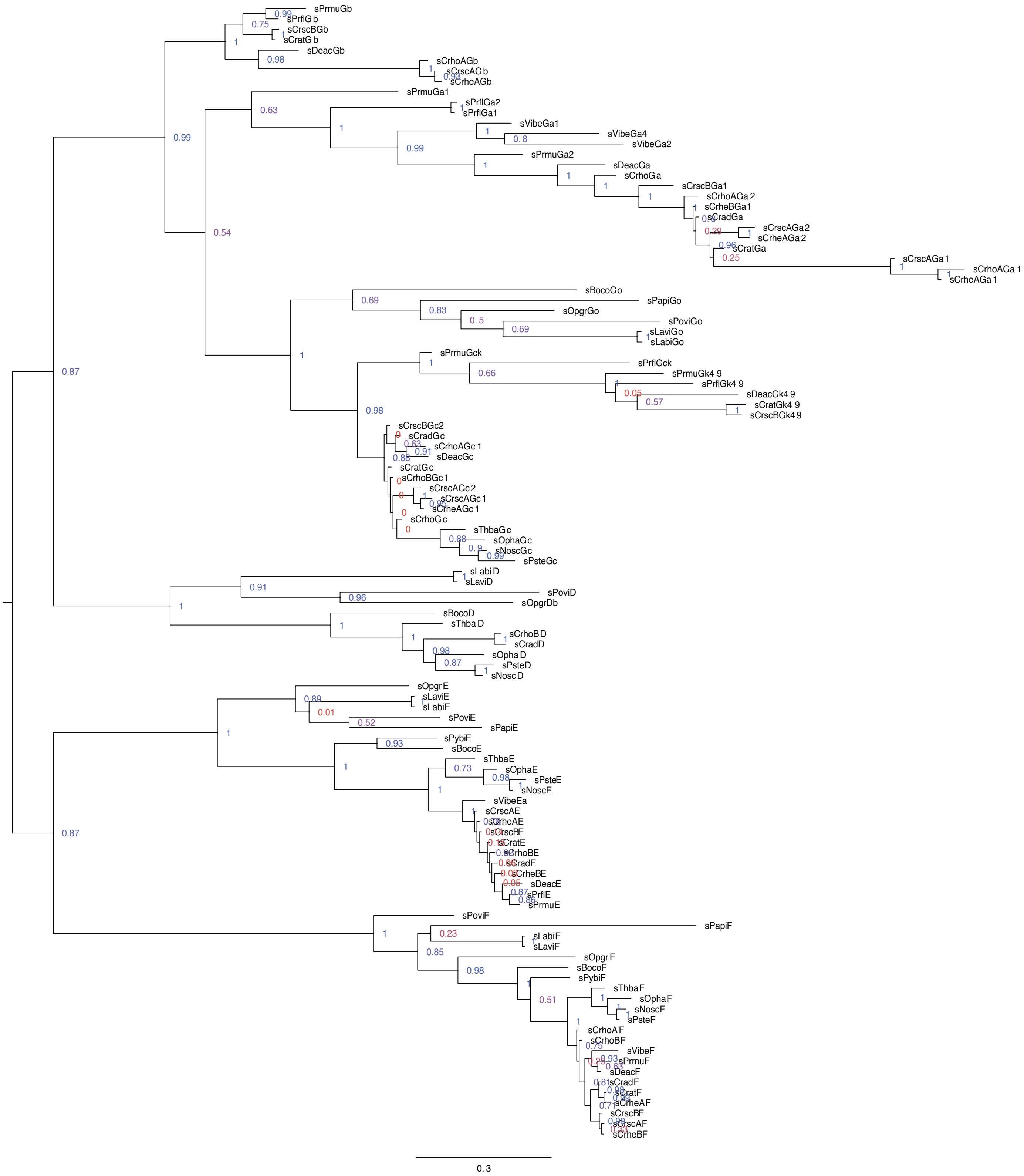

**Fig. 17. Phylogenetic tree of Pla2g2 genes of Squamata.** The tree is rerooted on the split between D-clade and EFC-clade.

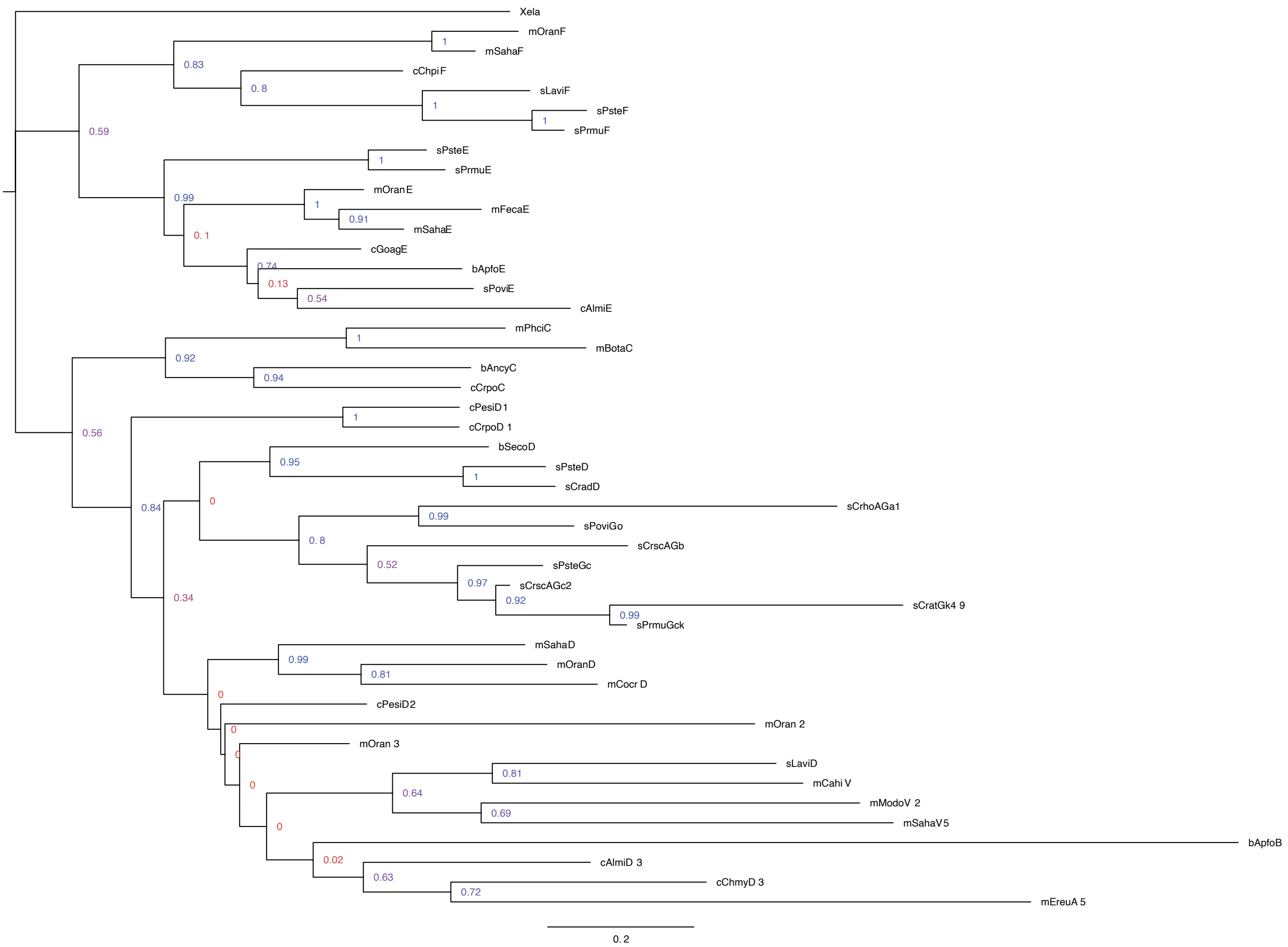

**Fig. 18. Phylogenetic tree of a single gene of each of the Pla2g2 clade.** The tree was not rerooted.

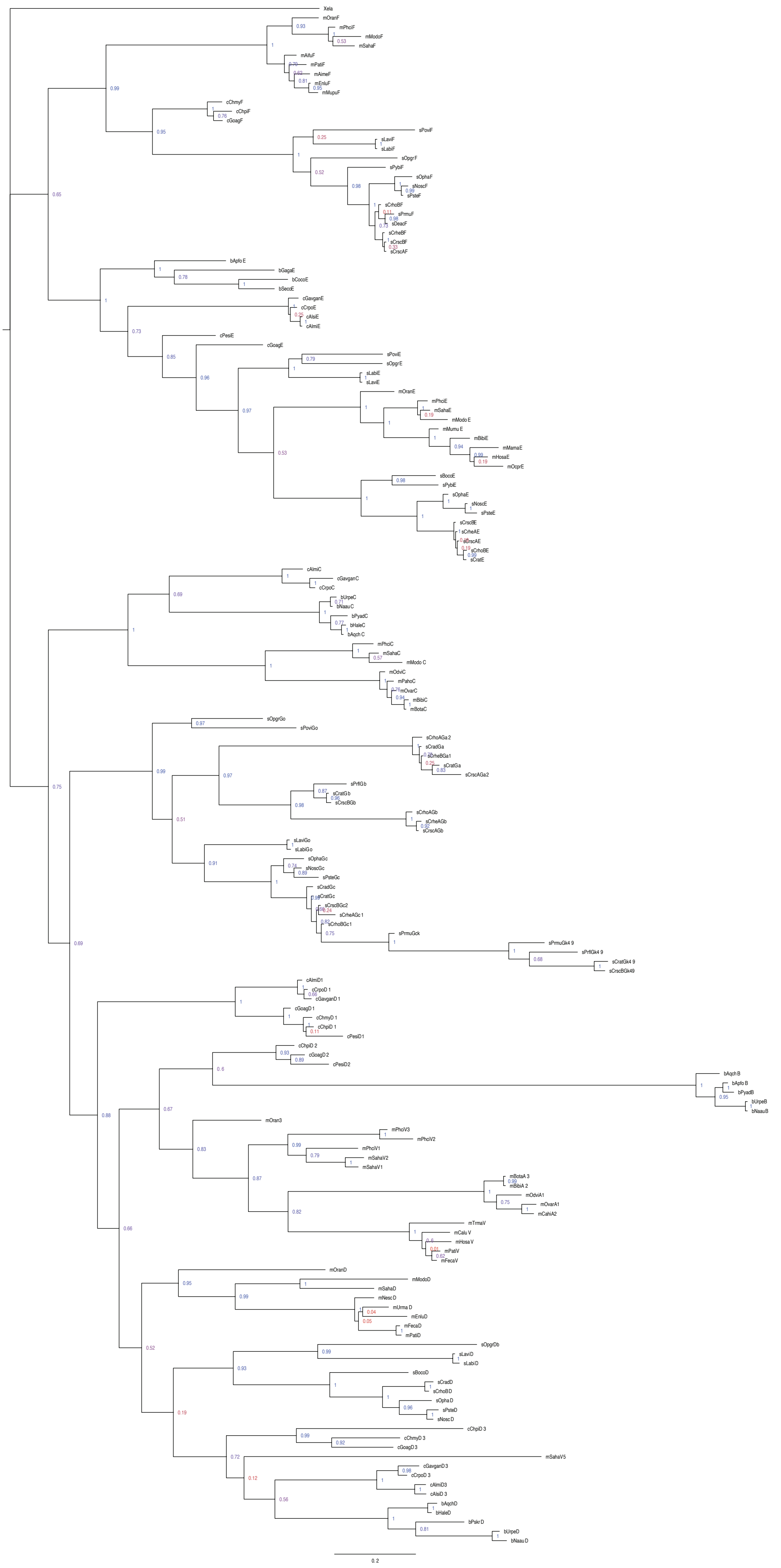

**Fig. 19. Phylogenetic tree of several (up to five where available) genes of each of the Pla2g2 clade.** The tree was not rerooted. Its larger topology is consistent with the tree in Fig.18. However most of the nodes have higher resolution and support value, as well as topology of inner nodes is different and more in accordance with all the other trees presented in this study.

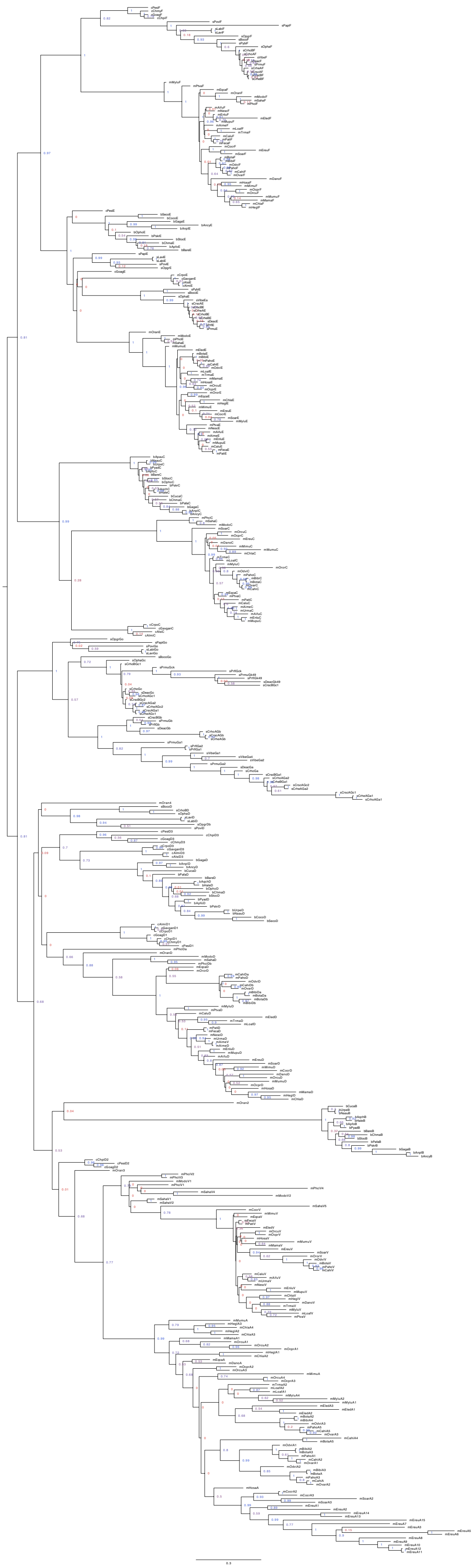

**Fig. 20. Phylogenetic tree of all Pla2g2 genes that were considered suitable for phylogenetic analysis.** The tree was rerooted on the split of EFC clade and D-clade.

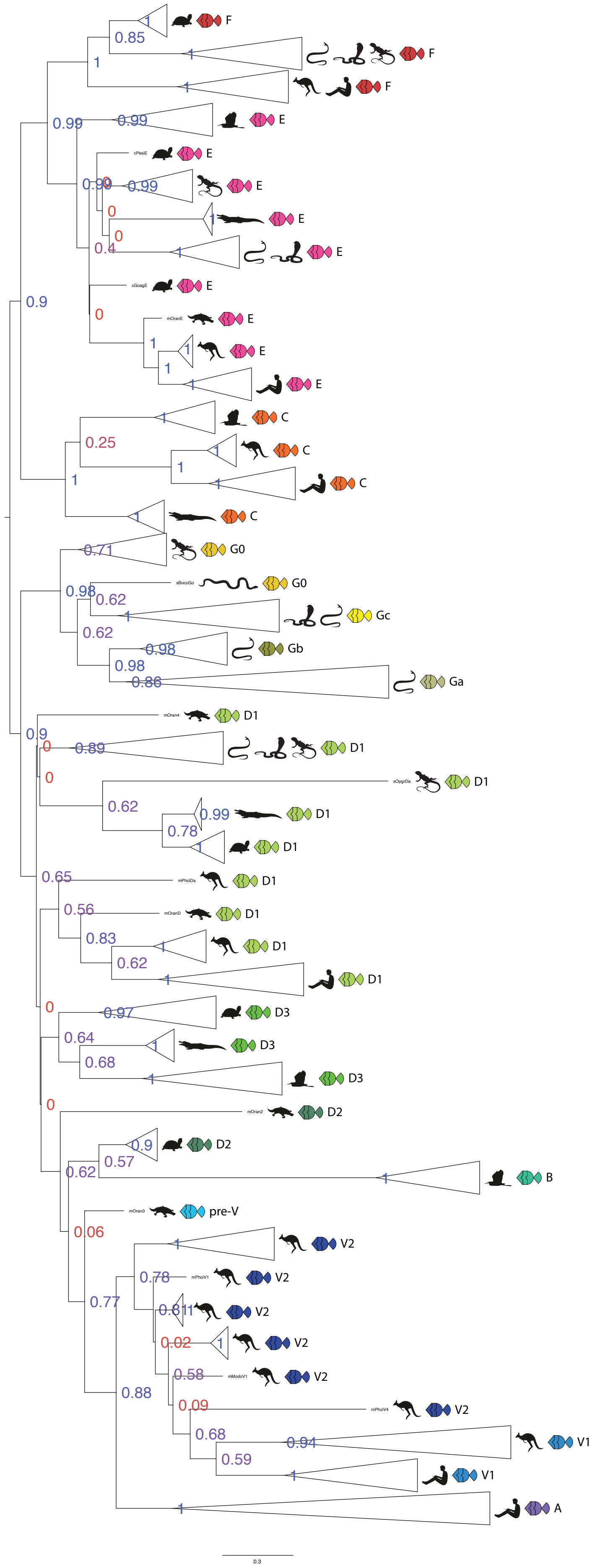

**Fig. 21. Phylogenetic tree of all Pla2g2 genes with each Pla2g2 clade collapsed.** This figure is based on a different tree than in Fig.20, it has lowered node supports but its topology is illustratively the same, The differences are in positions of marsupial g2V clades in respect to each other and some other intra-clade differences that are not important for the purposes of this study. The tree was rerooted on the split of EFC clade and D-clade.

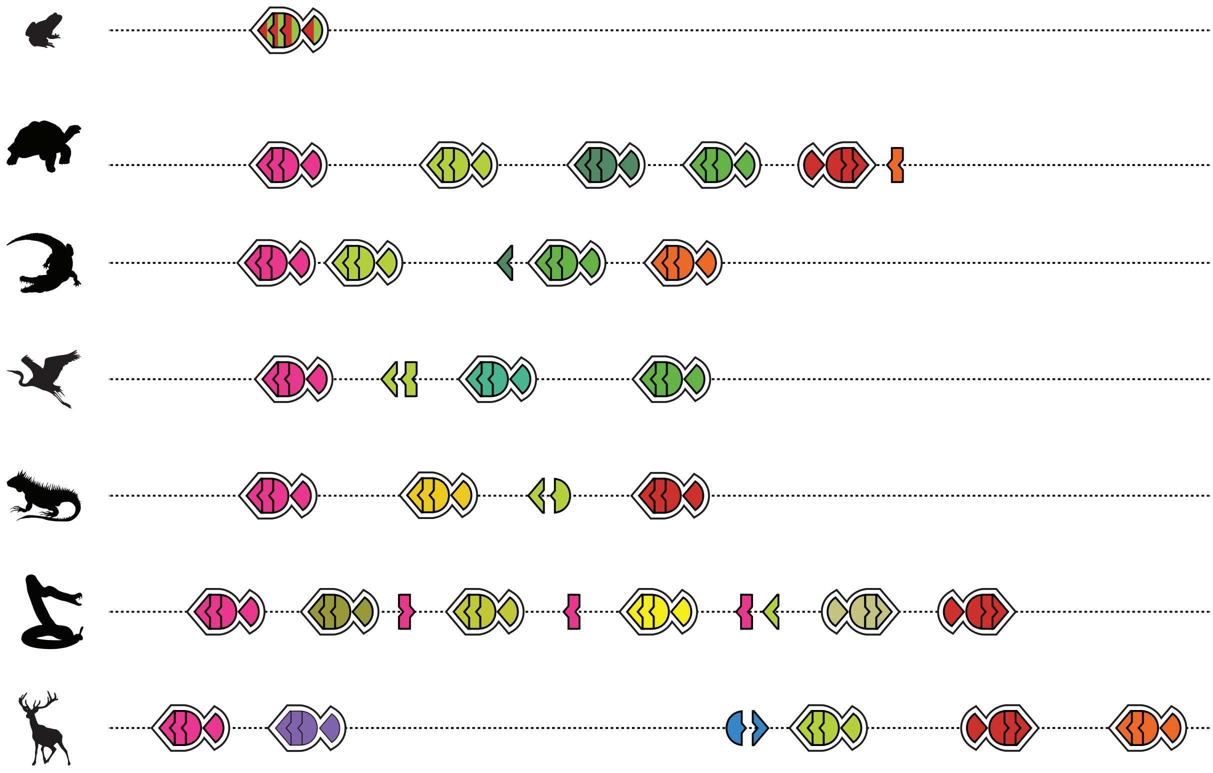

**Fig. 22. Step 5 of establishing homology between Pla2g2 genes: assigning gene homology based on phylogenetic analysis and genomic position.** Note that animal icons stand for a general representative of a group and not a particular species. And these genomic maps are for illustration only. For example, deer has 5 group 2 A genes and a functional group 5 gene, while a typical viper cluster has more venom genes than illustrated here.

#### *A note on nomenclature and the history of Pla2g2 research*

Despite the fact that research on Pla2s was initiated in the early 1900s (with studies of cobra venom and pancreatic juice), the nomenclature of this family wasn't formalized until the late 1990s, when many mammalian forms were cloned and studied. The earliest attempt at a systematic Pla2 nomenclature was that of Heinrikson et al. (Heinrikson, Krueger, and Keim 1977), who compared all available Pla2 sequences and, based on their structural features, proposed to lump mammalian pancreatic and elapid snake venom Pla2s together as Group 1 and viperid snake venom Pla2s as Group 2. Later, Joubert et al. (Joubert, Townshend, and Botes 1983) proposed to split Group 2 into g2A and g2B, the former of which included all known viperid sequences with the sole exclusion of a Bitis gabonica (gaboon viper) Pla2 which was classified as the single member of g2B. Later still, g2A was expanded to include mammalian synovial Pla2, when Davidson and Dennis (Davidson and Dennis 1990) made the first-ever Pla2 phylogeny using 40 protein sequences. Due to computational restrictions, they trimmed their dataset to include only one protein sequence per species, and their nomenclatural conclusions may have differed had their dataset included all sequences available at the time.

By the end of the 20th century all mammalian Pla2g2 subgroups had been discovered (Chen et al. 1994; Ishizaki et al. 1999; Valentin et al. 1999), and they received their names as a continuation of the g2A and g2B series: g2C, g2D, g2E, g2F. The only obvious exclusion was so-called "Group 5", that owed its special status to a reduced number of disulfide bonds (6 instead of 7) and the lack of knowledge about non-mammalian, non-squamate Pla2g2 sequences that routinely share this feature (Fig. 23, Fig. 13, SM6). Later it was discovered that "Group 5" is located within the chromosomal locus occupied by other Group 2 Pla2 genes in humans, but the nomenclatural distinction persisted.

At the same time, venom researchers started to use "gA" and "gB" to mean "acidic venom Pla2s" and "basic venom Pla2s" (cf. Whittington, Mason, and Rokyta 2018), which, given the historical grouping of viperid venom Pla2s and mammalian g2A was potentially confusing to researchers aiming to connect different kinds of Pla2s under a single system. To address this issue, Dowell et al. (Dowell et al. 2016) proposed to name all viperid venom Pla2s "g2G", with an additional distinction between acidic, basic and other distinct lineages within this subgroup.

Since our study has revealed more than 20 new Pla2g2 lineages (tripling in size the known number of subgroups), it has generated a need to resolve all conflicts within the nomenclature (ideally without creating additional confusion), while expanding it to include all Pla2g2s from the entire Vertebrata clade. In the interests of putting forth a system that takes into account the evolutionary relationships revealed in this study, we have taken the following steps:

- Extension of g2E, g2F and g2C to include all non-mammalian homologs that clearly clustered with their mammalian counterparts both in terms of phylogenetic relationship and chromosomal position.
- Expansion of clade g2D to include all Pla2s that cluster together with mammalian g2D but are not experiencing duplication or visible change of structure (unlike mammalian g2V/g2A, bird g2B or squamate g2G). However, we used indices to mark the deep evolutionary splits within the group (i.e. g2D1, g2D2 and g2D3).
- It should be noted that groupings within the "D-clade" highlight the differences between nomenclature and phylogeny. Our nomenclatural choices were principally motivated by the need to create a usable set of names for genes within this family and not by a requirement that these names conform to the demands of cladistics. For example, as the mammalian g2V, avian g2B, and squamate g2G clades are all derived from g2D2, they group phylogenetically with this gene to the exclusion of g2D1 and g2D3, i.e. g2D2 is more closely related to (e.g.) g2G than it is to g2D1.
- As the grouping of mammalian g2A and viperid venom Pla2s together has long been recognized as dubious (Six and Dennis 2000) and taking into account recent suggestions to label venom Pla2g2s as g2G (Dowell et al. 2016), we decided to use g2A to mean exclusively the uniquely eutherian Pla2g2 derived from Pla2g2D2 (see Fig. 14 through 20).
- Downgrading so-called "group 5" and placing it where it truly belongs, based on all available knowledge – as a part of group 2, thus labelling it as g2V1.
- Acknowledging the difference between those g2V that are present in both marsupials and placentals and those unique to marsupials, we decided to use g2V1 to mean the former and g2V2 to mean the latter. A unique Pla2g2 from platypus that seems to be basal to the entire g2V clade (g2V1, g2V2, g2A) thus received the name of pre-g2V.
- Because historical g2B, reserved solely for Bitis gabonica, has long been considered a misnomer (Six and Dennis 2000), and given the necessity to label a distinct clade of Pla2s from birds, the N-terminal region of which is dramatically different from all other Pla2g2 surveyed and is highly basic, we used g2B to label bird basic Pla2g2s.
- For the venom Pla2s, we have largely followed the nomenclature proposed by Dowell et al. only expanding it to include elapid Pla2g2s virtually indistinguishable from their viperid homologues, as well as non-venom (to the best of our knowledge) Pla2g2s from lizards, since they cluster together with g2G. The latter got the name of g2G0 to reflect their functionally incipient (i.e. pre-venom) state.

Thus, the chicken genes formerly known as "g2A" and "g5" were renamed to be g2D and g2C respectively, and mammalian "group 5" became g2V since it evolved from a group 2 precursor and in turn gave rise to g2A (Fig. 26). In addition, g2V has no distinct structural features that would justify its classification as a separate Pla2 group – the features previously used to describe it are in fact shared by several other g2 genes, which acquired it independently.

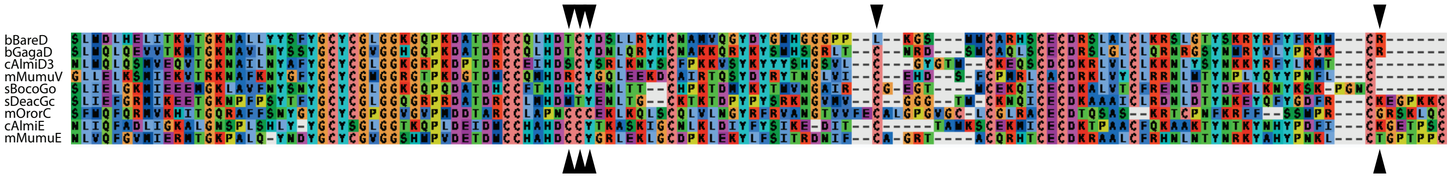

**Fig. 23. Convergent evolution of the cysteine arrangement formerly utilized as the defining characteristic of "Group 5" Pla2g2 in representatives of bird g2B and g2D, crocodile g2D3, snake g2G0 and mammalian g2V.**

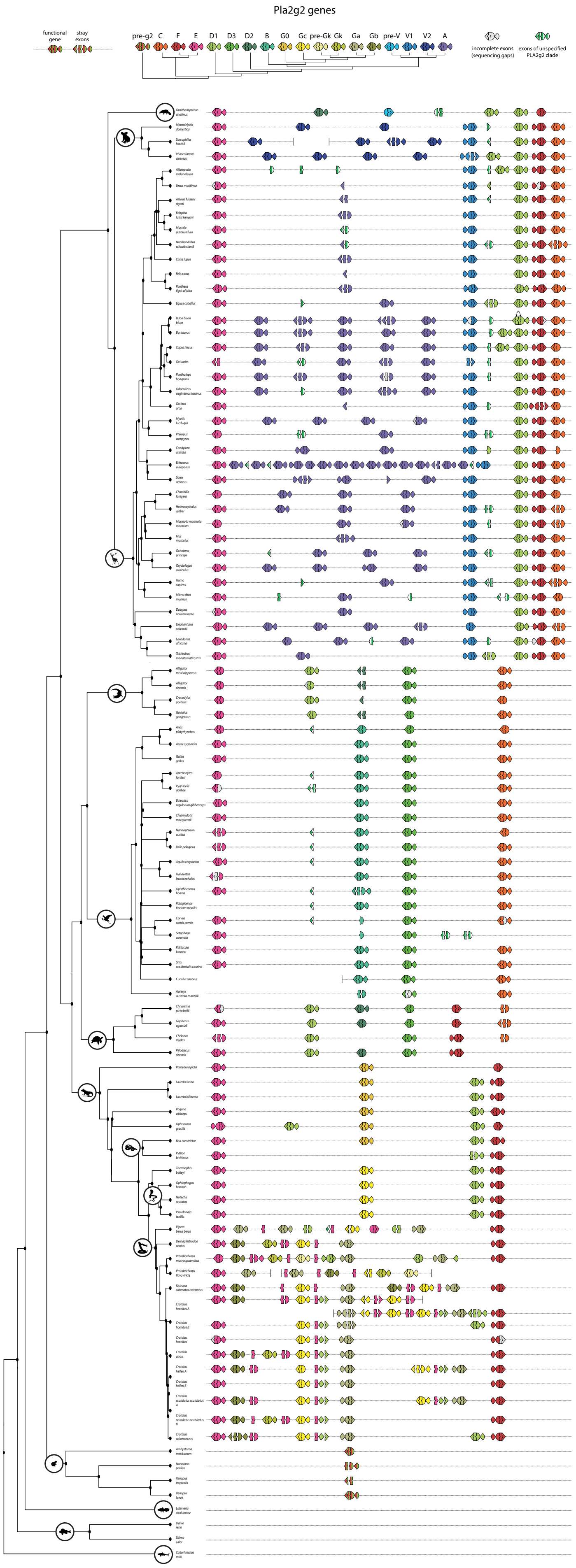

**Fig. 24. All exonic maps of Pla2g2 cluster that were made over the course of this study.** Phylogenetic tree is created with TimeTree online tool with closely related species substituting the ones actually used if needed.

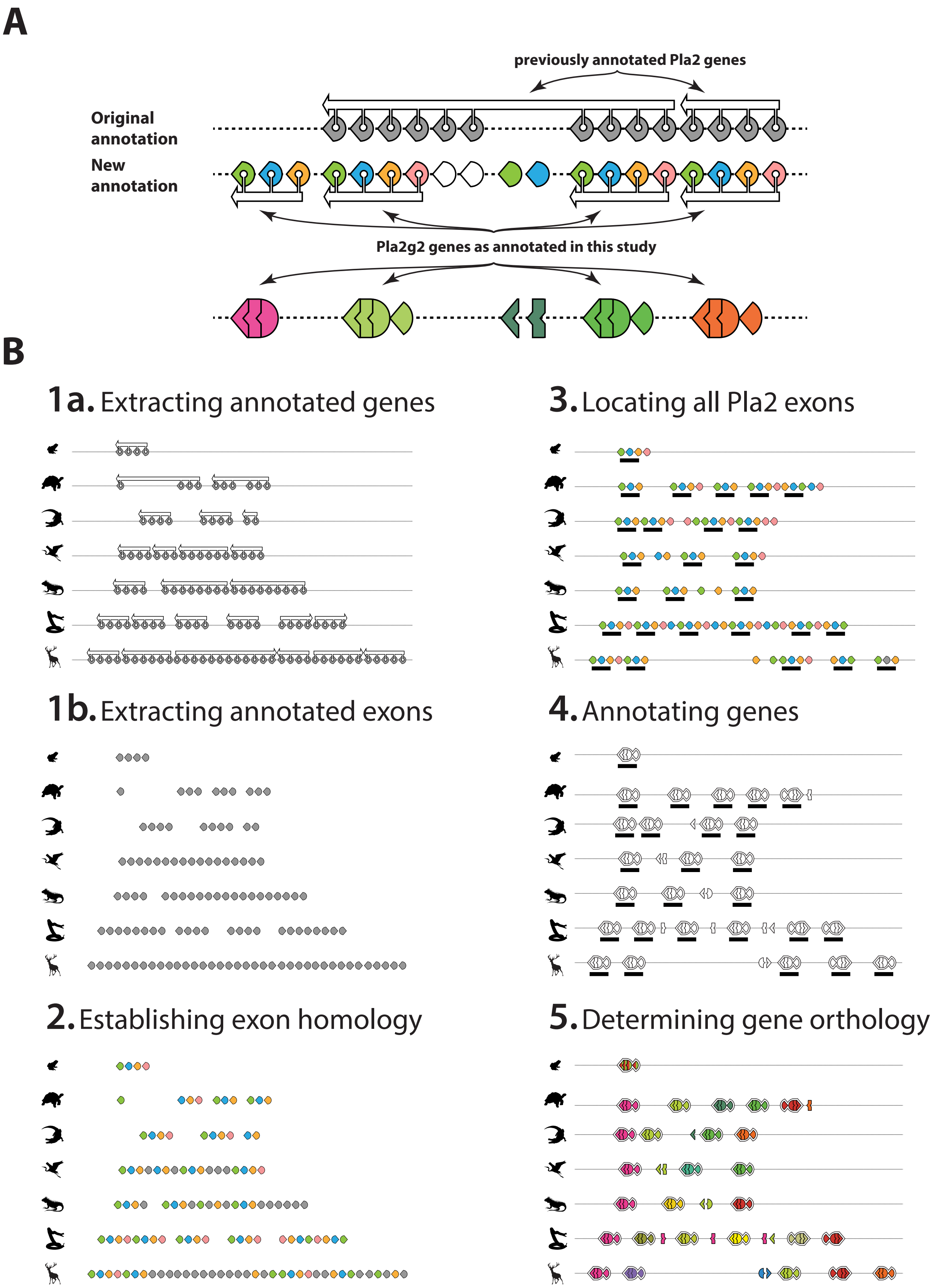

**Fig. 25.** ***Ab initio*** **annotation of Pla2g2 genes. A:** Side by side comparison of the approach employed in the present study and a published annotation (region from *Alligator mississippiensis* genome accessed last on 12.2018); **B:** Illustrative brief of our approach to gene annotation developed and implemented in this study.

#### *Selection analyses*

The entire dataset that was used for the construction of the phylogenetic trees was used for the analysis of selection. To detect signatures of natural selection and gauge the regime of selection dictating the evolution of various Pla2g2 lineages, site- and branch-site specific maximum likelihood models implemented in CodeML of the PAML (Phylogenetic Analysis by Maximum Likelihood) package were used (Yang 2007), and the omega parameter (ω), or the ratio of non-synonymous to synonymous substitutions, was estimated. To determine the statistical significance of the results obtained from nested models M7 (null model) and M8 (alternate model), the likelihood scores were compared with a likelihood ratio test (LRT). Amino acid sites under the influence of positive diversifying selection were identified using the Bayes Empirical Bayes (BEB) approach in M8 (Yang, Wong, and Nielsen 2005). Datamonkey webserver was used to assess the influence of episodic selection using Mixed Effect Model of Evolution (MEME) and the pervasive effects of diversifying and purifying selection using Fast Unconstrained Bayesian AppRoximation (FUBAR) analysis (Murrell et al. 2012, 2013). For assessing the nature of selection underpinning various Pla2g2 lineages, branch-site specific two ratio-model (Yang 1998; Yang and Nielsen 1998) was employed by marking the Pla2g2 lineage suspected to be evolving under positive selection as foreground (ω≥1, alternate model assuming positive selection), and by constraining others as background lineages (ω≤1, null model assuming negative selection or neutral evolution). The likelihood estimates of the null and alternate models were compared with an LRT for determining the significance. See Table 1 and 2 for results.

**Table 1. Pla2g2 evolutionary rate analyses.**

| Group | FUBAR[A] | MEME Sites[B] | PAML[C] (M8) |
|---|---|---|---|
| Birds | w>1[D]: 0 | 6 | 1 |
| | w<1[E]: 54 | | w: 0.33 |
| Crocodiles and turtles | w>1[D]: 0 | 5 | 0 |
| | w<1[E]: 38 | | w: 0.50 |
| Mammals | w>1[D]: 3 | 5 | 0 |
| | w<1[E]: 85 | | w: 0.32 |
| Otoconin-22-like | w>1[D]: 0 | 1 | 0 |
| | w<1[E]: 62 | | w: 0.14 |
| Squamates | w>1[D]: 1 | 7 | 0 |
| | w<1[E]: 66 | | w: 0.37 |
| Combined (five representatives) | w>1[D]: 0 | 2 | 0 |
| | w<1[E]: 86 | | w: 0.26 |

**A:** Fast Unconstrained Bayesian AppRoximation (FUBAR), **B:** Sites identified under the influence of episodic diversifying selection (0.05 significance) by the Mixed Effects Model Evolution (MEME), **C:** Positively selected sites detected by the Bayes Empirical Bayes approach implemented in M8. Sites detected at PP≥0.95, **D:** Sites experiencing pervasive diversifying selection at the posterior probability ≥0.9 (FUBAR), **E:** Sites experiencing pervasive purifying selection at the posterior probability ≥0.9 (FUBAR), ω: mean dN/dS

**Table 2. Branch-site model estimations of the evolutionary rate of Pla2g2 groups.**

| Foreground | ω | Likelihood (l) | Sites with ω > 1 |
|---|---|---|---|
| Pla2g2 – A and V | **2.42**** | -25820.68 | 1 |
| Pla2g2B | 1.00 | -25796.35 | 8 |
| Pla2g2C | 1.00 | -25810.92 | 2 |
| Pla2g2 – E and F | 1.00 | -25796.68 | 1 |
| Pla2g2G | **2.34*** | -25798.20 | 5 |
| Pla2g2G of snakes | **2.75**** | -25792.63 | 5 |
| Pla2g2Gc | **2.17** | -25823.20 | 0 |

**ω:** mean dN/dS ('*' – p<0.05, '**' – p<0.01)

#### *Graphical summary of syntenic information gathered in this study that led to our theories on the evolution of the cluster*

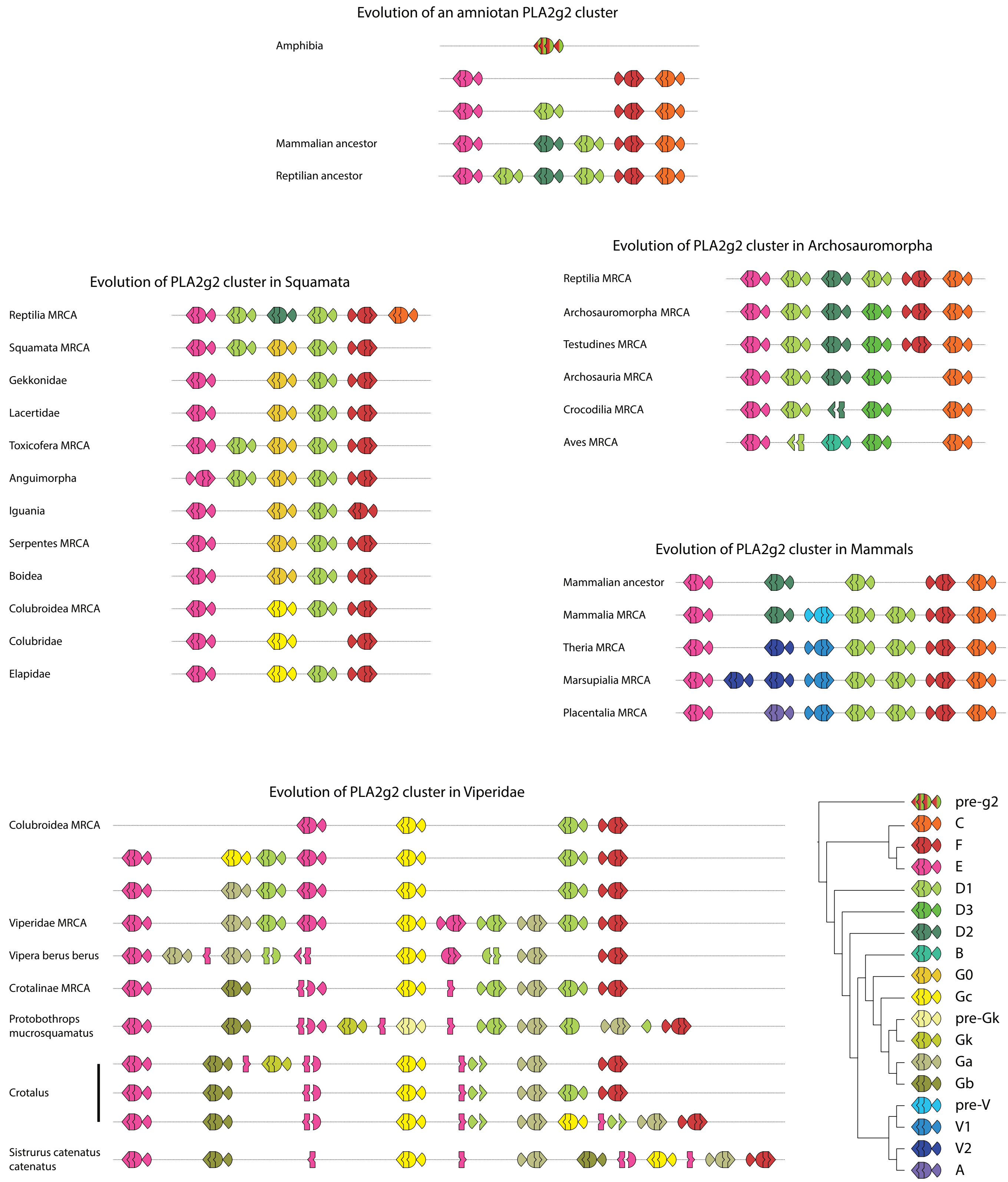

**Fig. 26. Reconstructions of evolutionary paths that are hypothesized to have led to the existence of the Pla2g2 cluster in present day animals.** Inference is based on the exon maps as they are presented in Fig. 24 with gene relationships derived from the sum of phylogenetic analyses (as presented in Fig. 14 – 20). The number of steps could have been more, since we favored lower number of steps over higher in our deduction.

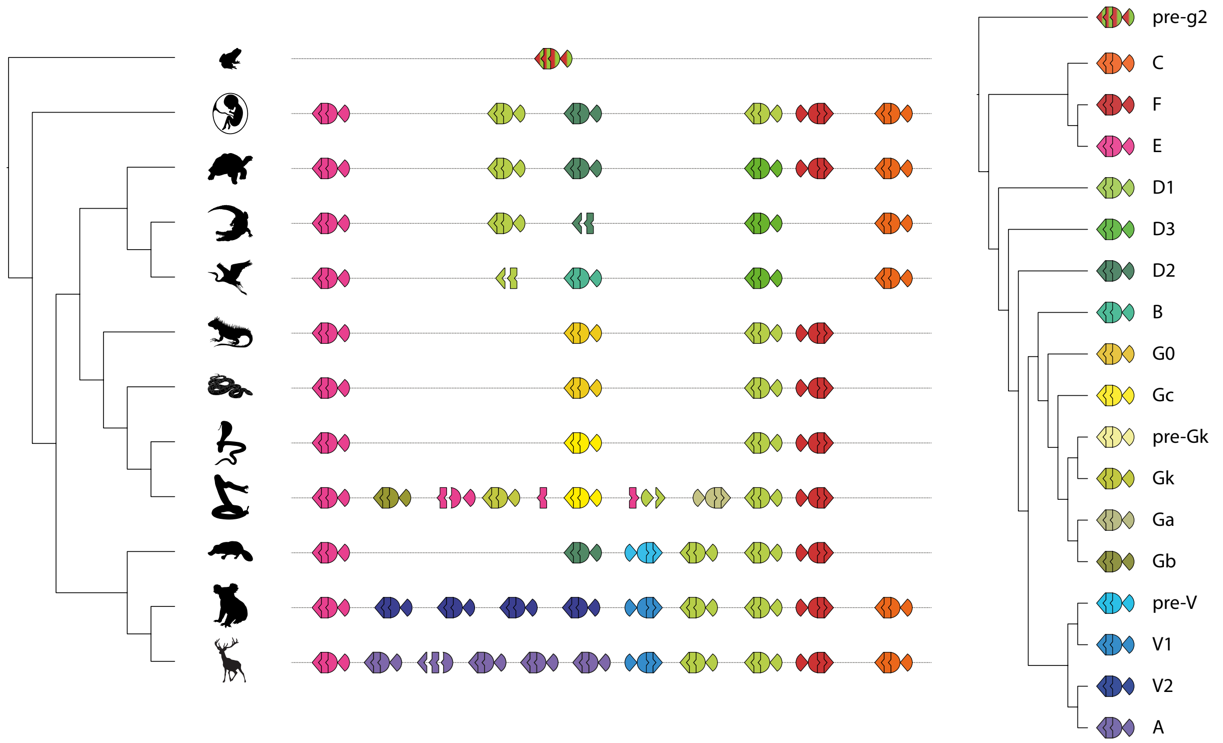

**Fig. 27. Schematic representations of Pla2g2 clusters across the Vertebrata and relationships between the genes as recovered in this study.**

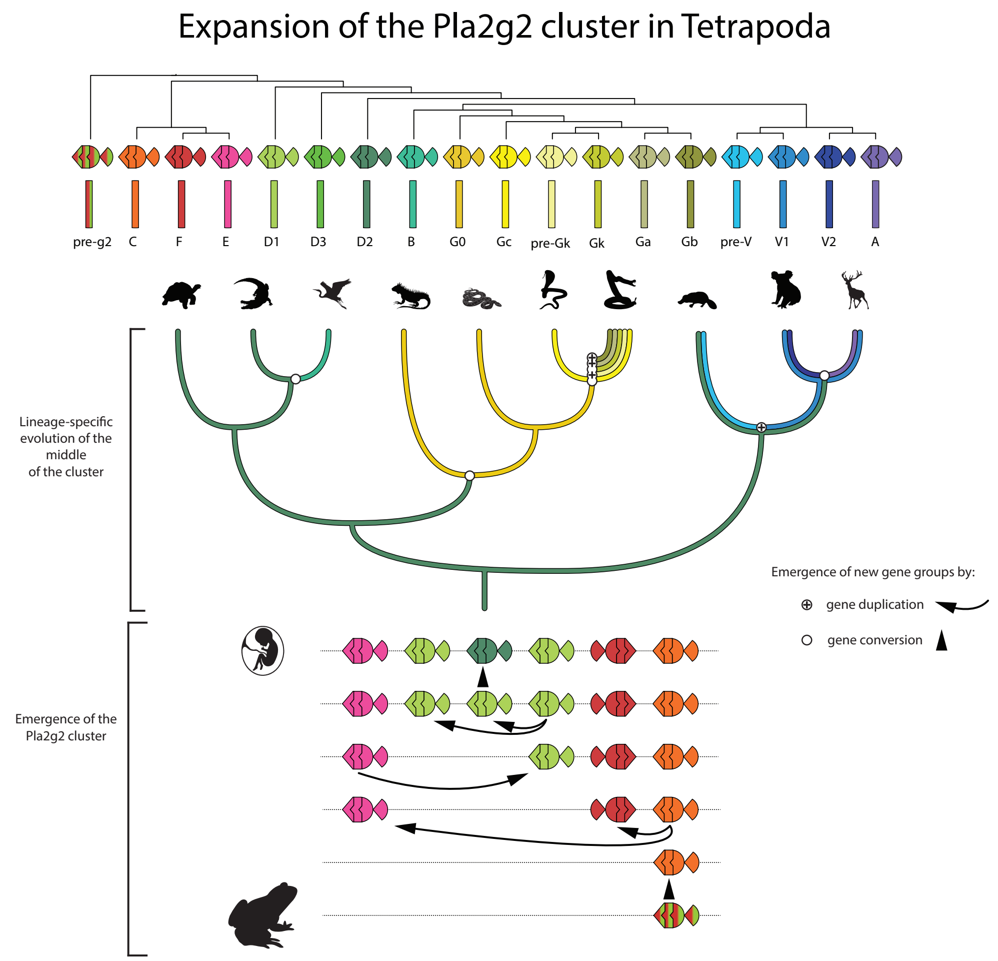

**Fig. 28. Evolution of the Pla2g2 cluster in Tetrapoda. A:** Cladogram depicting relationships among Pla2g2 groups and key to color-coding. **B:** Inferred expansion of the cluster following the split of amniotes from amphibians. Note that since the inferred amniote MRCA all subsequent expansion has taken place at the Pla2g2 D2 locus.

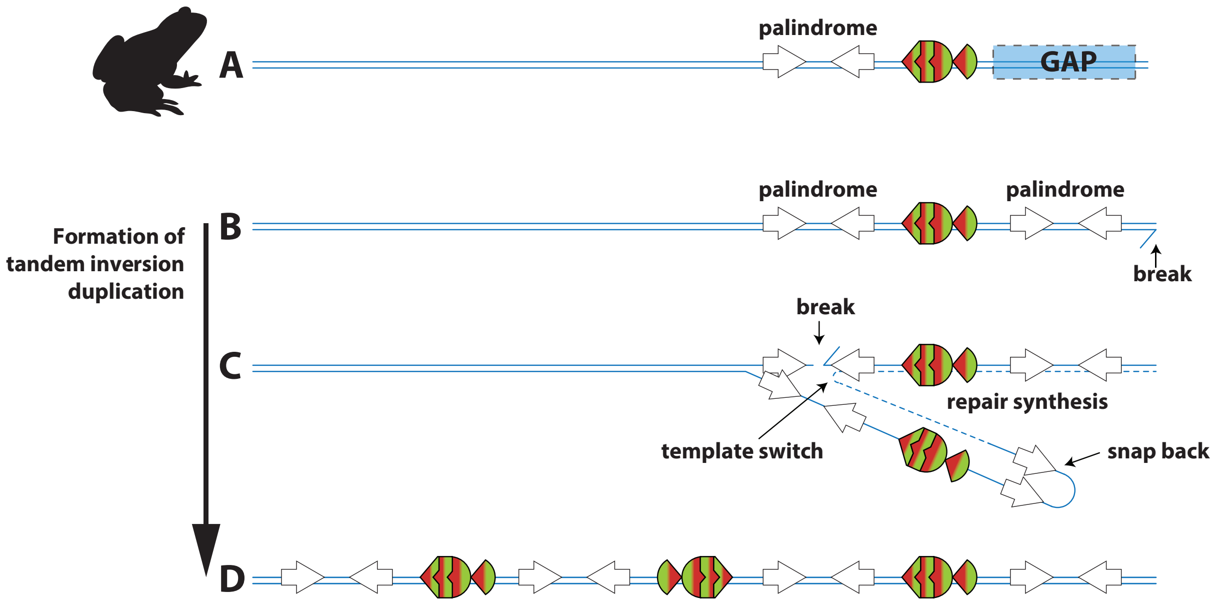

Fig. 29. Illustration of the tandem-inversion duplication mechanism hypothesized to have created the EFC clade of Pla2g2 (adapted from (Reams and Roth 2015). A: Illustration of the extant condition of Xenopus laevis and X. tropicalis, where GAP indicates a sequencing gap present in published genomes of both species. B and C: following a DNA break the presence of flanking palindromic sequences (parts of the transposon "Kolobok T2") results in template switching leading to the incidental multiplication of genes during the repair process. D: The final result is 3 genes, with two copies in the same orientation as the parent gene and one reversed.

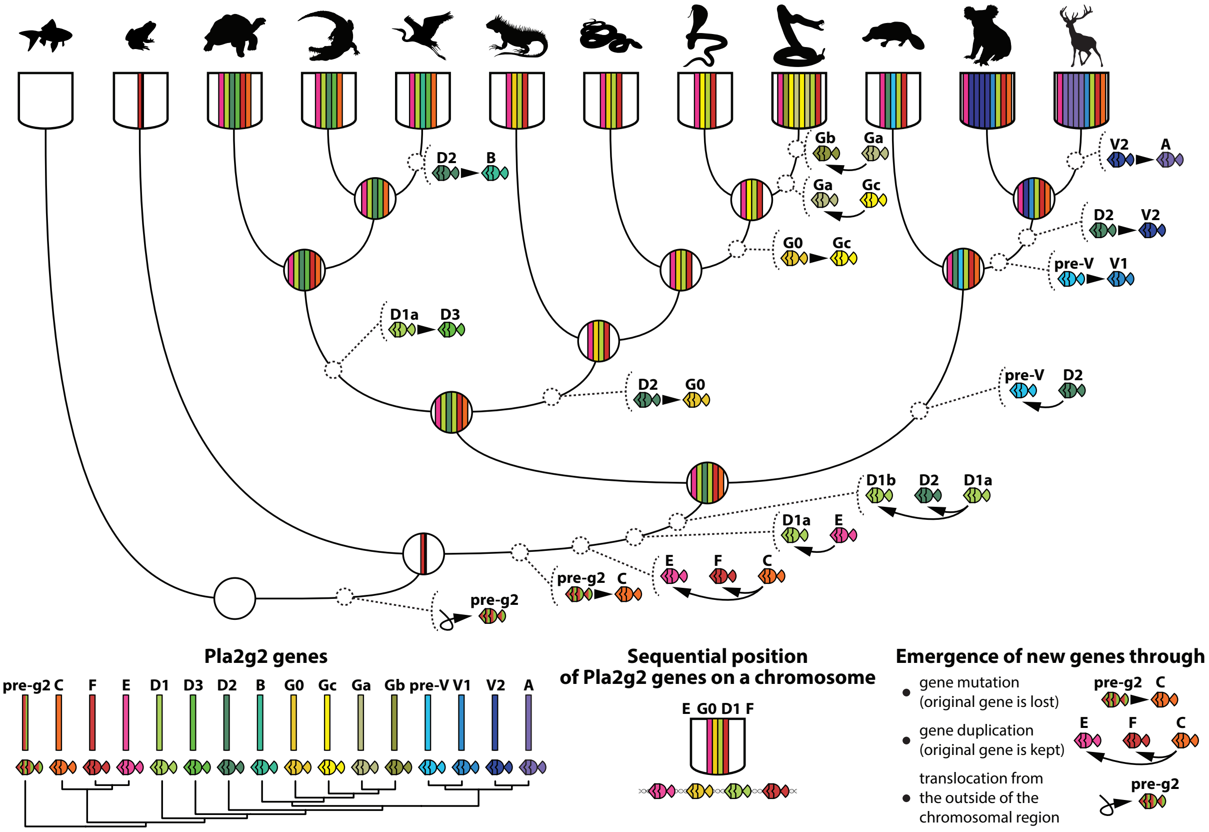

Fig. 30. Graphical summary of our theory on the evolution of the cluster with key events highlighted.

#### *Evolution of the mammalian Pla2g2A clade*

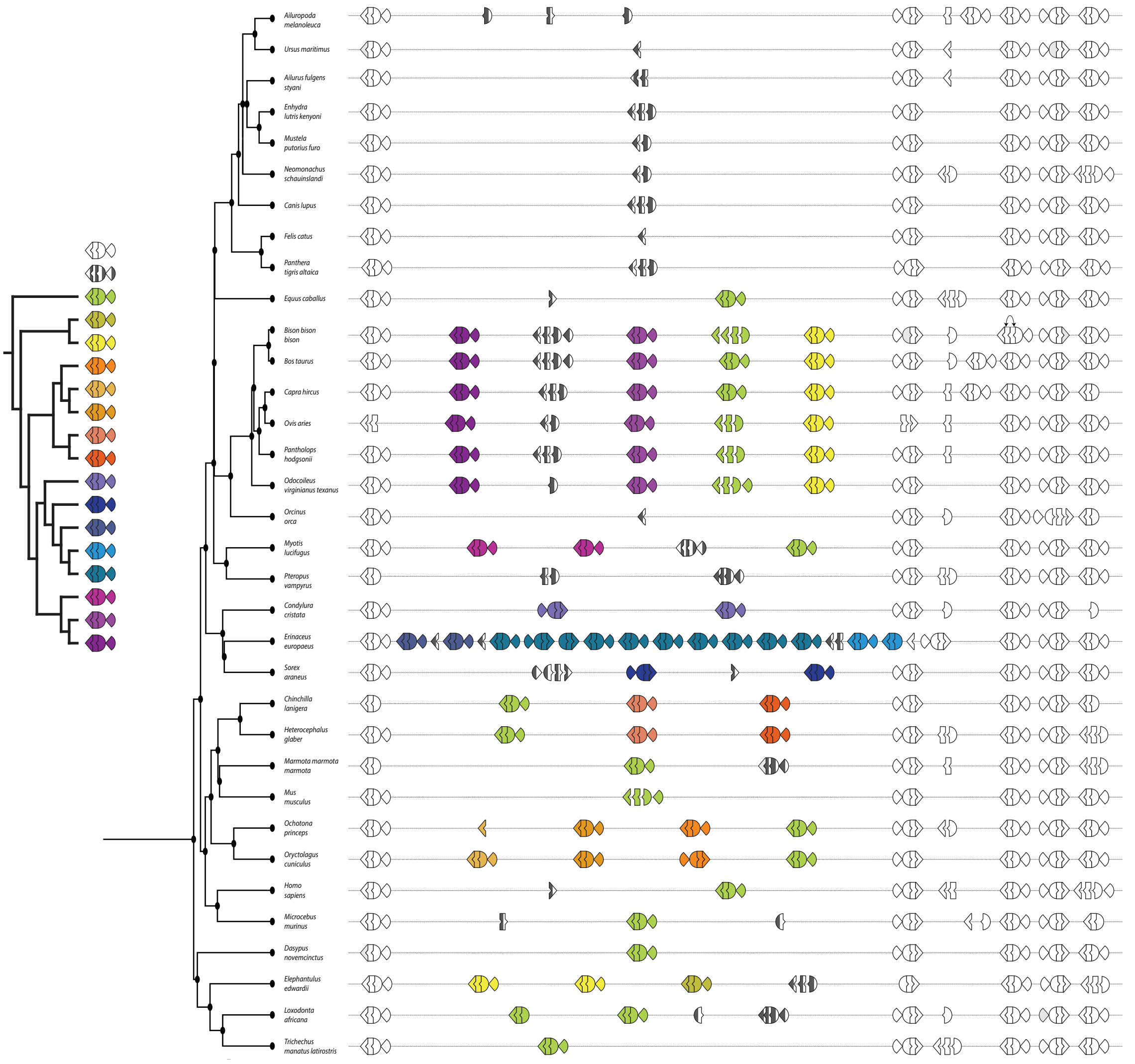

SM1-12. Phylogenetic and syntenic relationships within the placental Pla2g2A clade. Pla2g2 of other clades (g2E, g2V, g2D, g2F, g2C) are colored white. Genes and exons of g2A clade that cannot be unambiguously assigned a groupping within the g2A radiation are colored in black and white stripes. A shared group that is present in most of placental lineages and the one we consider ancestral to all g2A is colored bright green.

**Fig. 31. Phylogenetic and syntenic relationships within placental Pla2g2A clade.** Pla2g2 of clades other than A (i.e. g2E, g2V, g2D, g2F, g2C) are colored white to increase the clarity of the complexities of relationships within the g2A clade. Genes and exons of g2A that could not be unambiguously assigned a grouping within the g2A radiation are colored in black and white stripes. The shared group that is present in most of placental lineages and the one we consider ancestral to all other g2A is colored lime green.

#### *Evolution of the reptilian Pla2g2G clade*

The branching pattern within the g2G clade is convoluted and presents a challenge to the formulation of a rational nomenclature for these genes that both accurately reflects the intraclade relationships and preserves previous taxonomic systems (e.g. Dowell et al. 2016). The deepest split within this clade is either that between lizard forms (herein labelled "G0") and the various snake forms (supplementary material SM4) or between lizard and Boa copies and colubroid snake copies (supplementary material SM1-9; see also SM5 for consensus primary structures). The second node marks the split between viperid snake only acidic and basic forms and all other snake-specific copies. These latter genes include the plesiotypic form from Boa constrictor (again labelled "G0" in recognition of its plesiotypy and similarity to lizard forms, with which it clusters in some analyses e.g. supplementary material SM1-9) and the various forms of "Gc". "Gck" and "Gk" are structurally derived forms of "Gc", with the "Gck" transitional between "Gc" and "Gk". We note the challenges of delineating a "new gene" on the basis of structural change and again flag the important caveat that structural and functional changes are not synonymous (Jackson and Fry, 2016) – here we have based our classification of G0 and Gc on the lineage-specific (Colubroidea only) appearance of the Gc clade, which clusters together to the exclusion of the plesiotypic G0 forms found in all other squamate reptiles (Fig. 6; supplementary material SM1-9). As with all such classification schemes, this should be considered schematic – our subsequent analysis would be unchanged were we to group G0 and Gc together under the same name.

Figure 32 illustrates the effect of the acquisition of a novel function in venom on subsequent lineage-specific gene family expansion. Although viper snake and elapid snake g2Gc are homologous and 94% similar at the sequence level, only viperid snakes utilise the gene product utilized as a toxin, and further expansion has occurred only within this lineage.

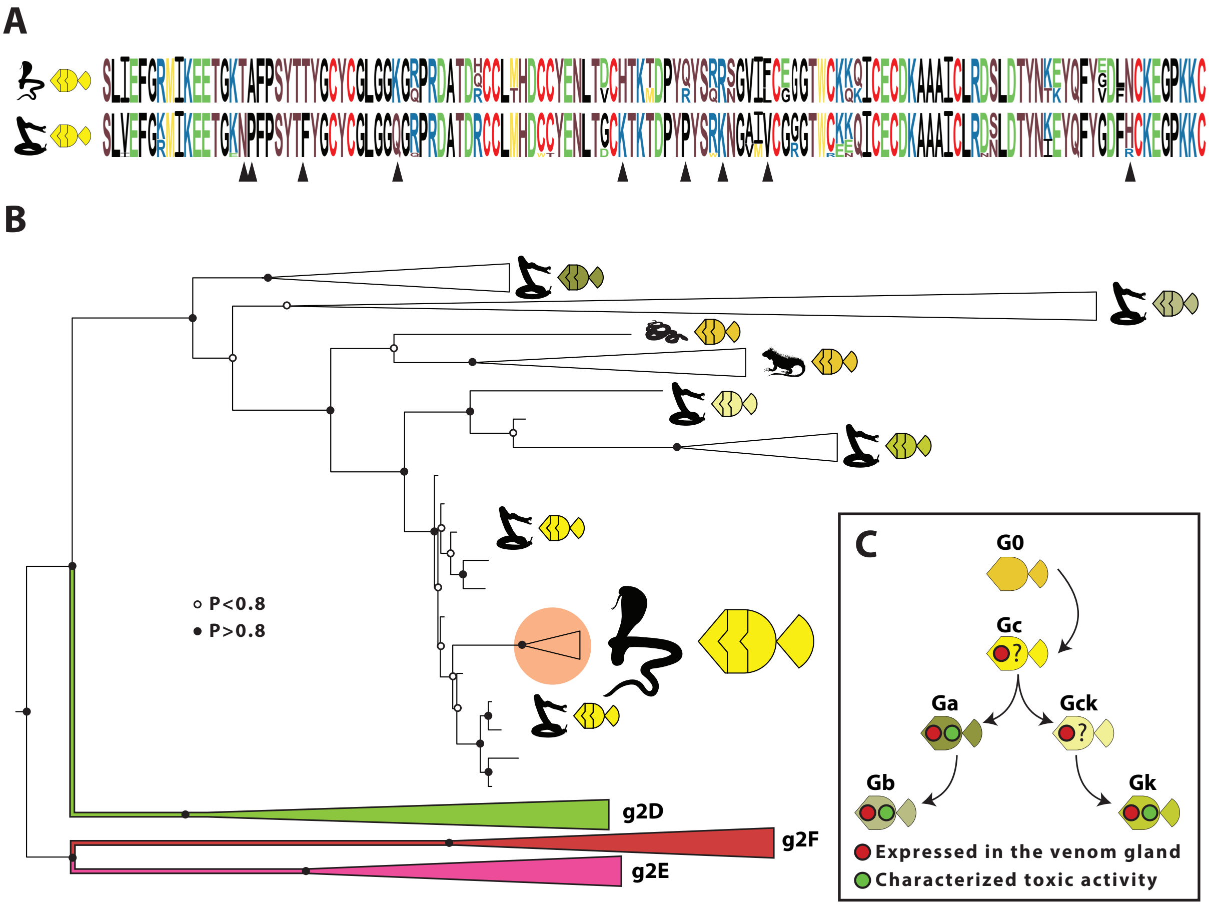

**Fig. 32. Comparison of viperid (utilized as a toxin) and elapid (not utilized as a toxin) snake Pla2g2G. A:** Consensus sequences of elapid g2Gc and viperid g2Gc, with differences between them indicated by black triangles. Overall, the sequences are 94% similar. The size of the letters is proportional to the probability of a member of the lineage having it; **B:** Phylogenetic tree of squamate Pla2g2 genes, with the orange circle highlighting the nesting of elapid g2Gc sequences within viperid venom sequences, a consequence of the extreme similarity of these forms; **C:** Reconstruction of the evolution of the Gc group, including current evidence of toxic activity and expression in viperid snake venom glands. Based on all available evidence, single-copy gene g2Gc became expressed in the venom gland prior to its multiplication that created all the modern viper venom Pla2g2 genes and therefore should be regarded as the original viper venom Pla2g2.

#### *Final notes*

While this manuscript was in preparation, we used the approach developed for this study to successfully locate and annotate several gene families albeit at a far less ambitious scale. The results were published in three papers: vertebrate NAD glycohydrolases (Koludarov and Aird, 2019), mammalian kallikreins (Casewell et al. 2019) and Indian cobra three finger toxins (Suryamohan et al. 2020).

#### *References*

Aberer AJ, Kobert K, Stamatakis A. 2014. ExaBayes: massively parallel bayesian tree inference for the whole-genome era. Mol. Biol. Evol. 31:2553–2556.

Aird SD, Arora J, Barua A, Qiu L, Terada K, Mikheyev AS. 2017. Population Genomic Analysis of a Pitviper Reveals Microevolutionary Forces Underlying Venom Chemistry. Genome Biol. Evol. 9:2640–2649.

Bradnam KR, Fass JN, Alexandrov A, Baranay P, Bechner M, Birol I, Boisvert S, Chapman JA, Chapuis G, Chikhi R, et al. 2013. Assemblathon 2: evaluating de novo methods of genome assembly in three vertebrate species. Gigascience 2:10.

Casewell NR, Petras D, Card DC, Suranse V, Mychajliw AM, Richards D, Koludarov I, Albulescu LO, Slagboom J, Hempel BF, Ngum NM, et al. 2019. Solenodon genome reveals convergent evolution of venom in eulipotyphlan mammals. Proc. Natl. Acad. Sci. U. S. A. 116: 25745-25755.

Chen J, Engle SJ, Seilhamer JJ, Tischfield JA. 1994. Cloning and recombinant expression of a novel human low molecular weight Ca (2+)-dependent phospholipase A2. J. Biol. Chem. 269:2365–2368.

Chijiwa T, Hamai S, Tsubouchi S, Ogawa T, Deshimaru M, Oda-Ueda N, Hattori S, Kihara H, Tsunasawa S, Ohno M. 2003. Interisland mutation of a novel phospholipase A2 from Trimeresurus flavoviridis venom and evolution of Crotalinae group II phospholipases A2. J. Mol. Evol. 57:546–554.

Davidson FF, Dennis EA. 1990. Evolutionary relationships and implications for the regulation of phospholipase A2 from snake venom to human secreted forms. J. Mol. Evol. 31:228–238.

Dowell NL, Giorgianni MW, Kassner VA, Selegue JE, Sanchez EE, Carroll SB. 2016. The Deep Origin and Recent Loss of Venom Toxin Genes in Rattlesnakes. Curr. Biol. 26:2434–2445.

Dowell NL, Giorgianni MW, Griffin S, Kassner VA, Selegue JE, Sanchez EE, Carroll SB. 2018. Extremely divergent haplotypes in two toxin gene complexes encode alternative venom types within rattlesnake species. Curr. Biol. 28:1016-1026.

Heinrikson RL, Krueger ET, Keim PS. 1977. Amino acid sequence of phospholipase A2-alpha from the venom of Crotalus adamanteus. A new classification of phospholipases A2 based upon structural determinants. J. Biol. Chem. 252:4913–4921.

Ishizaki J, Suzuki N, Higashino K-I, Yokota Y, Ono T, Kawamoto K, Fujii N, Arita H, Hanasaki K. 1999. Cloning and Characterization of Novel Mouse and Human Secretory Phospholipase A2s. J. Biol. Chem. 274:24973–24979.

Jackson TNW, Fry BG. 2016. A tricky trait: Applying the fruits of the "function debate" in the philosophy of biology to the "venom debate" in the science of toxinology. Toxins, 8(9):263.

Joubert FJ, Townshend GS, Botes DP. 1983. Snake Venoms. Purification, some Properties of Two Phospholipases A2(CM-I and CM-II) and the Amino-Acid Sequence of CM-II from Bitis nasicornis (Horned Adder) Venom. Hoppe-Seyler´s Zeitschrift für physiologische Chemie 364:1717–1726.

Katoh K, Standley DM. 2013. MAFFT multiple sequence alignment software version 7: improvements in performance and usability. Mol. Biol. Evol. 30:772–780.

Koludarov I, Aird SD, 2019. Snake venom NAD glycohydrolases: primary structures, genomic location, and gene structure. PeerJ 7:e6154.

Kumar S, Stecher G, Suleski M, and Hedges SB, 2017. TimeTree: a resource for timelines, timetrees, and divergence times. Molecular Biology and Evolution 34: 1812-1819.

Larsson A. 2014. AliView: a fast and lightweight alignment viewer and editor for large datasets. Bioinformatics 30:3276–3278.

Murrell B, Moola S, Mabona A, Weighill T, Sheward D, Kosakovsky Pond SL, Scheffler K. 2013. FUBAR: a fast, unconstrained bayesian approximation for inferring selection. Mol. Biol. Evol. 30:1196–1205.

Murrell B, Wertheim JO, Moola S, Weighill T, Scheffler K, Kosakovsky Pond SL. 2012. Detecting individual sites subject to episodic diversifying selection. PLoS Genet. 8:e1002764.

Ogawa T, Oda-Ueda N, Hisata K, Nakamura H, Chijiwa T, Hattori S, Isomoto A, Yugeta H, Yamasaki S, Fukumaki Y, Ohno M. 2019. Alternative mRNA Splicing in Three Venom Families Underlying a Possible Production of Divergent Venom Proteins of the Habu Snake, Protobothrops flavoviridis. Toxins. 11:581.

Six DA, Dennis EA. 2000. The expanding superfamily of phospholipase A2 enzymes: classification and characterization. Biochimica et Biophysica Acta (BBA) – Molecular and Cell Biology of Lipids 1488:1–19.

Suryamohan K, Krishnankutty SP, Guillory J, Jevit M, Schröder MS, Wu M, Kuriakose B, Mathew OK, Perumal RC, Koludarov I, Goldstein LD, et al. 2020. The Indian cobra reference genome and transcriptome enables comprehensive identification of venom toxins. Nat Genet doi:10.1038/s41588-019-0559-8

Valentin E, Ghomashchi F, Gelb MH, Lazdunski M, Lambeau G. 1999. On the diversity of secreted phospholipases A(2). Cloning, tissue distribution, and functional expression of two novel mouse group II enzymes. J. Biol. Chem. 274:31195–31202.

Venturini L, Caim S, Kaithakottil GG, Mapleson DL, Swarbreck D. 2018. Leveraging multiple transcriptome assembly methods for improved gene structure annotation. GigaScience. 7, 8.

Wang Y, Liu J, Huang BO, Xu YM, Li J, Huang LF, Lin J, Zhang J, Min QH, Yang WM, Wang XZ. 2015. Mechanism of alternative splicing and its regulation. Biomedical reports. 3:152-158.

Whittington AC, Mason AJ, Rokyta DR. 2018. A

Single Mutation Unlocks Cascading Exaptations in the Origin of a Potent Pitviper Neurotoxin. Mol. Biol. Evol. 35:887–898.

Yang Z. 1998. Likelihood ratio tests for detecting positive selection and application to primate lysozyme evolution. Mol. Biol. Evol. 15:568–573.

Yang Z. 2007. PAML 4: phylogenetic analysis by maximum likelihood. Mol. Biol. Evol. 24:1586–1591.

Yang Z, Nielsen R. 1998. Synonymous and nonsynonymous rate variation in nuclear genes of mammals. J. Mol. Evol. 46:409–418.

Yang Z, Wong WSW, Nielsen R. 2005. Bayes empirical bayes inference of amino acid sites under positive selection. Mol. Biol. Evol. 22:1107–1118.
